# *Bacteroides*-driven metabolic remodelling suppresses *Clostridioides difficile* toxin expression in mixed biofilm communities

**DOI:** 10.1101/2025.09.03.674004

**Authors:** Kavana Kalea Bywater-Brenna, Simran Kaur Aulakh, Kiran Raosaheb Patil, Niranjan Nagarajan, Meera Unnikrishnan

## Abstract

*Clostridioides difficile* is a major cause of hospital-associated diarrhoea worldwide. The intricate interactions between *C. difficile* and the resident gut microbiota play a crucial role in determining the outcome of *C. difficile* infection (CDI), although the molecular mechanisms underlying many *C. difficile*-commensal interactions are not understood. Here we show that selected *Bacteroides* species can inhibit *C. difficile* growth within mixed biofilms. A transcriptomic analysis of *C. difficile-Bacteroides* biofilms showed significant metabolic shifts, with distinct changes in carbohydrate and amino acid metabolism and, interestingly, a downregulation of *C. difficile* toxin gene expression. A significant reduction in *C. difficile* toxin production was evident in *C. difficile*-*Bacteroides* cocultures, irrespective of the extent of *C. difficile* growth inhibition. Notably, Stickland fermentation of proline, which is known to repress toxin synthesis, was upregulated in *C. difficile*, while proline synthesis was induced in the cocultured species *B. vulgatus* and *B. dorei*. Furthermore, upregulation of proline reductase pathways and consequent toxin repression were evident within a synthetic 9-species gut commensal biofilm community containing multiple *Bacteroides* spp. Thus, leveraging multiomics approaches, we demonstrate a potential cross-feeding mechanism where proline produced by *B. dorei* and *B. vulgatus* is utilised by *C. difficile* through Stickland fermentation to drive toxin repression. Our study reveals a new mechanism of microbiota-mediated control of a key virulence factor involved in *C. difficile* pathogenesis while enabling pathogen co-existence within a polymicrobial commensal community.

**Importance:** *C. difficile* infection, characterised by severe diarrhoea and colitis, has a significant impact on healthcare settings globally due to the high rates of recurrence. CDI is closely associated with the gut microbiota status and the use of antibiotics, yet the mechanistic basis of interactions between the causative bacterium *C. difficile* and individual gut commensal species remains poorly defined. Here, we demonstrate inhibitory effects of *Bacteroides* species on *C. difficile* through nutrient competition and a cross-feeding mechanism between these abundant gut commensals and this pathogen which blocks expression of key *C. difficile* virulence factors. Our findings offer insights into the effective design of microbiota consortia to prevent and treat CDI.

## Introduction

*Clostridioides difficile* is a spore-forming obligate anaerobic pathogen and aetiological cause of *C. difficile* infection (CDI), which can lead to severe recurrent diarrhoea, pseudomembranous colitis and in rare instances, toxin megacolon^1,2^. CDI is strongly associated with antibiotic therapy and with distinct changes in the gut microbiota composition^3–5^. It has become clear that the healthy gut microbiota plays a critical role in preventing *C. difficile* spore germination and proliferation, with recent studies providing new insights into mechanisms underlying colonisation resistance against CDI^6–9^. Several *C. difficile* virulence factors have been identified, with the toxins A, B and CDT being shown to play a key role in the epithelial damage and the subsequent inflammatory responses elicited in CDI^10^. The impact of the gut microbiota on *C. difficile* virulence factors, however, remains unclear.

The *Bacteroides* genus is one of the most abundant genera in the human microbiome, constituting approximately 25-50% of the overall gut microbiota^11,12^. CDI is typically characterised by the significant underrepresentation of Bacteroidetes^13^, with antibiotic treatment commonly resulting in its near-to-complete eradication^14,15^. Successful faecal microbiota transplantations (FMTs) focus largely around the restoration of Bacteroidetes, particularly among the *Bacteroides* genus^14^. *Bacteroides* abundance has generally been negatively correlated with that of *C. difficile* during CDI^16,17^. However, a few studies have reported a positive association with *Bacteroides* in CDI-positive individuals, and the cause of this discrepancy is unclear^18–21^. *Bacteroides* are known for their glycan-degrading capabilities and production of antimicrobial proteins, and play a significant role in shaping the community structure in the gut by modulating nutrient availability and influencing microbial growth^22–24^. The reduced prevalence of *Bacteroides* in cases of CDI suggests a critical role for these bacteria in preventing *C. difficile* colonisation, although specific mechanisms through which they protect from CDI are not well understood.

Several *Bacteroides* species populate the gut, but there have been limited studies on specific *Bacteroides*-*C. difficile* interactions. A recent screen highlighted that many *Bacteroides* species displayed inhibitory effects against *C. difficile in vitro*^25^. However, not all associations appear inhibitory; *C. difficile* has been recently shown to form symbiotic biofilms with *B. thetaiotaomicron* during antibiotic exposure, promoting its persistence^26^. The underlying mechanisms of these antagonistic or synergistic interactions are not known.

While there have been several commensal-pathogen coculture studies to date, the vast majority of these have been conducted using planktonic cultures^25,27,28^. However, it is important to consider that gut bacteria are not strictly planktonic, and can attach to the colonic mucosa to form sessile biofilm communities^29^. *C. difficile* strains have been reported to form biofilms both *in vitro* and *in vivo*^30–33^, and have been shown to associate with commensals and microbiota communities^34–36^; microbiota communities containing *C. difficile* are also thought to act as reservoirs for recurrent infection^34^.

In this study we report *Bacteroides* species-specific inhibition of *C. difficile* within mixed biofilm communities. We further investigate *Bacteroides* spp. and *C. difficile* interactions combining conventional microbiology and multiomics approaches, demonstrating distinct changes in metabolic pathways that may drive reduction in *C. difficile* growth. We also find that multiple *Bacteroides* species can suppress *C. difficile* toxin production. We demonstrate that *Bacteroides* spp. produce proline, which promotes Stickland fermentation of proline in *C. difficile*, inhibiting toxin production. Finally, we highlight how Stickland fermentation contributes to the formation of stabilised but toxin-neutralised polymicrobial association of *C. difficile* within a synthetic commensal community.

## Results

### *C. difficile* growth is inhibited by specific *Bacteroides* species

*Bacteroides* spp. have been negatively associated with CDI^16^, and we previously demonstrated inhibitory effects by *Bacteroides* spp. on *C. difficile* growth^37–39^. We first sought to determine interactions of a range of *Bacteroides* species with the highly virulent ribotype 027 *C. difficile* R20291, when cocultured as biofilms.

Dual species *C. difficile-Bacteroides* biofilms cultured in Schaedler anaerobe broth (SAB+), which supports growth of both *C. difficile* and *Bacteroides* species (Fig S1A-D), were tracked over 48 h by measuring bacterial counts and biofilm biomass. Culturing *C. difficile* alongside *B. dorei*, *B. ovatus*, and *B. thetaiotaomicron* in coculture biofilms, showed a significant reduction in *C. difficile* bacterial numbers (Fig. 1A-C). The most potent inhibition of *C. difficile* was elicited in the *B. dorei/C. difficile* coculture at 24 h, which showed a striking ∼3,000,000-fold drop in *C. difficile* number, and in *B. ovatus/C. difficile* and B. *thetaiotaomicron/C. difficile* cocultures, with a 339- and 193-fold reduction of *C. difficile* respectively (Fig. 1A-C). While *C. difficile* cultured with either *B. vulgatus,* or *B. fragilis* had slight, but statistically non-significant detrimental effects, *B. finegoldii* had a very minor effect on *C. difficile* growth at 24 h (Fig. 1D-F).

**Figure 1.**
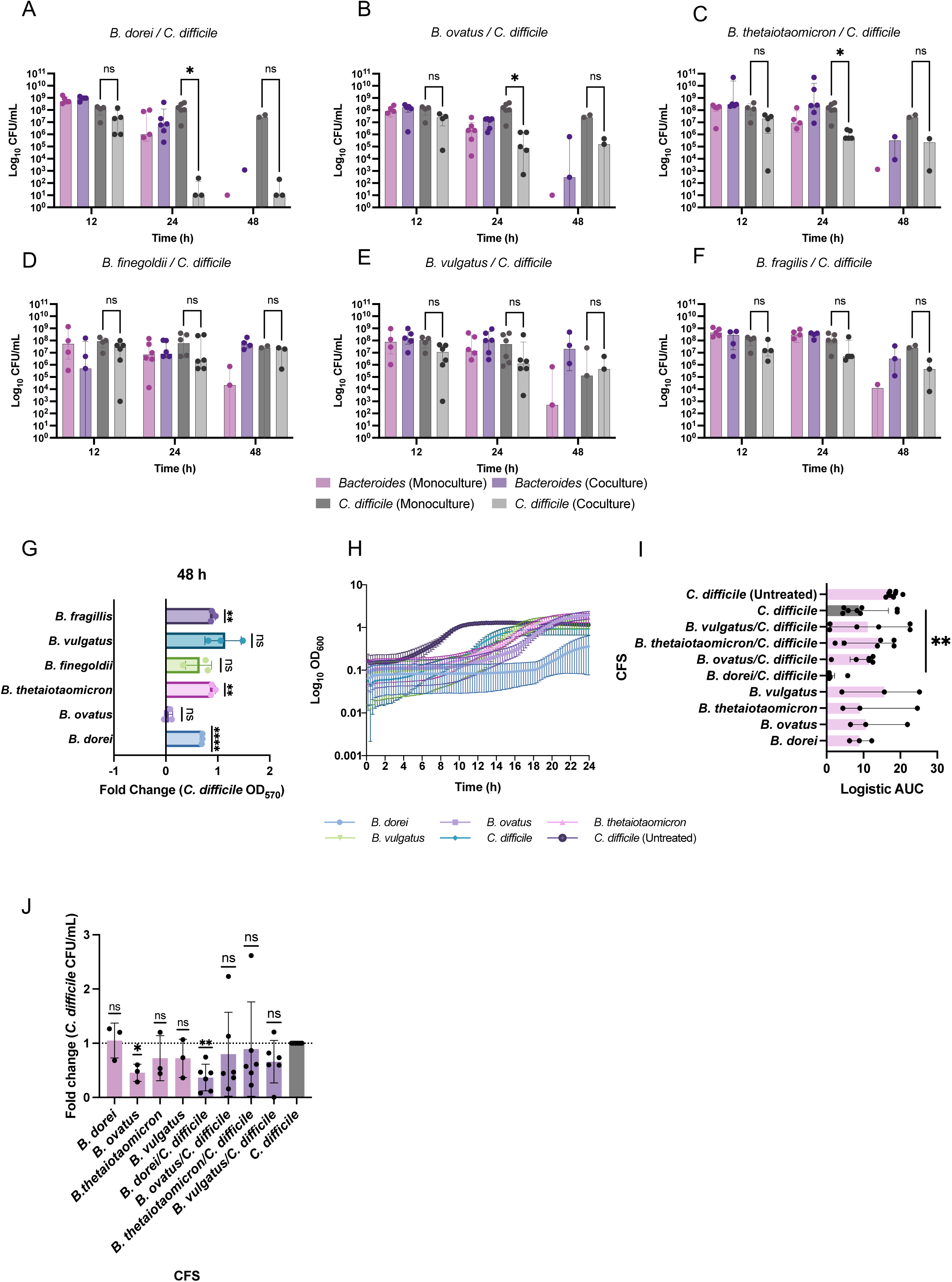
*C. difficile* inhibition by selected *Bacteroides* species. CFUs from monoculture and coculture biofilms were enumerated at 12, 24 and 48 h under anaerobic conditions for *C. difficile* cocultured with **A**) *B. dorei* **B**) *B. ovatus*, **C**) *B. thetaiotaomicron*, **D**) *B. finegoldii*, **E**) *B. vulgatus,* and **F**) *B. fragilis*. Panels display the median plus interquartile range. Data represents a minimum of 3 biological repeats. Multiple Mann– Whitney tests were performed with Holm–Šídák correction for multiple comparisons. **G)** Biomass of coculture biofilms from 48 h stained with crystal violet were quantified by measuring OD_570_ and normalised to the biomass of a *C. difficile* monoculture biofilm from 48 h. Fold changes are presented, with mean ± SD. N = 3. A one-sample t-test was performed against a hypothetical value of 1. **H)** *C. difficile* growth curves spiked with 50% of CFS derived from different coculture 24 h biofilms. A CFS control from a 24 h *C. difficile* monoculture biofilm was used. An untreated control *C. difficile* cultured in the absence of any CFS was run in parallel. Data is displayed as mean ± SEM. N = 6. **I)** Logistic AUCs were calculated based on growth curve data. Data represents a minimum of 3 biological repeats. Graph displays the median plus the interquartile range. Separate Mann-Whitney tests were performed to compare each CFS AUC with the *C. difficile* CFS AUC. ** indicates *P* = 0.0027. Data represents a minimum of three biological repeats. **J**) Fold change of CFU/mL from biofilms grown in 50% CFS was calculated, standardised to CFU/mL enumerated from 24 h *C. difficile* biofilms grown in 50% CFS derived from a control *C. difficile* 24 h biofilm. A Wilcoxon singed-rank test or one sample t-test was performed, from a hypothetical mean of 1: * indicates *P* = 0.0270, ** indicates *P* = 0.0015. Data represents a minimum of three biological repeats, showing the mean ± SD. Statistical denotations are represented by: ns (not significant) *P* > 0.05, * *P* < 0.05, ** *P* < 0.01, **** *P* < 0.0001.

Biofilm biomass measurements across time showed that certain *Bacteroides* species, *B. ovatus, B. thetaiotaomicron,* and *B. finegoldii* formed significantly less biofilm biomass, compared to *C. difficile* or cocultures at early timepoints (12 h) (Fig. S2A,B). A similar trend of increased biofilm biomass was seen for most cocultures compared to the monocultures of *Bacteroides* at 24 h, or compared to *C. difficile* monocultures at 48 h (Fig. S2B). Fold changes in biomass, normalised to the monoculture *C. difficile* biofilms showed enhanced biofilm formation in the presence of *Bacteroides* species at 48 h including the cocultures that showed decreased *C. difficile* numbers, but this was not observed at earlier timepoints (Fig. 1G, Fig. S2C). This may indicate that there is increased production of biofilm matrix components which may further impact *C. difficile* survival.

To study *C. difficile* strain specificity of this *Bacteroides*-mediated inhibition, we tested the *C. difficile* strain 630. A significant reduction in *C. difficile* numbers was still observed in biofilms cocultured with *B. dorei* and *B. ovatus* (Fig. S3). Although there was a trend in *C. difficile* reduction when *C. difficile* was cocultured alongside the other *Bacteroides* species, the inhibitory effects were not significant (Fig. S3A). All strains showed similar growth rates when cultured in SAB+ (Fig. S3B,C).

To investigate if *Bacteroides*-mediated inhibitory effects on *C. difficile* were also observed in planktonic culture, planktonic mono- and cocultures were assessed. A similar inhibition of *C. difficile* in coculture was observed, particularly at 24 h (Fig. S4). As in a biofilm milieu, *B. dorei*, *B. ovatus and B. thetaiotaomicron* elicited the most striking inhibition of *C. difficile* and lower *C. difficile* numbers were recovered when cultured with *B. vulgatus* or *B. fragilis* at 24 h (Fig. S4B).

To probe whether there was a secreted factor responsible for the inhibitory effects on *C. difficile*, cell free supernatants (CFS) were harvested from mono and coculture 24 h biofilms and added to *C. difficile* bacterial inoculums, prior to setting up planktonic or biofilm cultures. While monoculture CFS, including the *B. dorei* monoculture CFS, had no notable impact on *C. difficile* planktonic growth (Fig. 1I, Fig. S1E), *C. difficile* spiked in with a *B. dorei/C. difficile* CFS displayed overall poor planktonic growth kinetics compared to the control, with a severely delayed lag phase (Fig. 1H,I); 50% CFS derived from *B. dorei/C. difficile* coculture biofilms elicited a significant 53% reduction in *C. difficile* recovery from 24 h biofilms (Fig. 1J). These data suggest that the presence of both bacterial partners in the biofilm may be required to stimulate production of a secreted inhibitory factor. Although CFS had varying impact on *C. difficile* bacterial numbers within biofilms, there was negligible impact on overall biofilm biomass (Fig. S2D). Furthermore, mass spectrometry analysis comparing CFS from *B. dorei* and *C. difficile* monoculture and coculture biofilms, showed minimal changes in the *B. dorei* proteome (Fig. S5A). In fact, the majority of observed differences were attributed to alterations in the *C. difficile* proteome in coculture, with enrichment of genes involved in fructose metabolism (Fig. S5B). Interestingly, in the coculture there was increased abundance of a *B. dorei* β-D-fructan fructohydrolase, an enzyme responsible for hydrolysing fructans into fructose (Fig. S5C).

### A dual species RNAseq reveals metabolic shifts in *C. difficile*-*Bacteroides* biofilms

To investigate potential mechanisms underlying inhibition we studied the transcriptomes of *C. difficile* and *Bacteroides* spp. cultured as biofilms. RNA was extracted from *C. difficile* cocultured for 24 h with either *B. dorei* or *B. vulgatus*, which display different levels of inhibition of *C. difficile* (Fig. 1), and was processed and sequenced as described in the Methods. RNAseq analysis revealed distinct transcriptional shifts in *C. difficile* expression in the different cocultures relative to the monoculture (Fig. 2A, Fig. S6A). Of the significantly differentially expressed genes, 44 were commonly upregulated in the two cocultures, and 156 were commonly downregulated (Fig. 2B). Although there was a downregulation of common pathways by *C. difficile* in response to the *Bacteroides* partner, overall, a species-specific *C. difficile* transcriptional response was observed (Fig. 2B). A complete list of significant differentially expressed genes can be found in Table S3.

**Figure 2.**
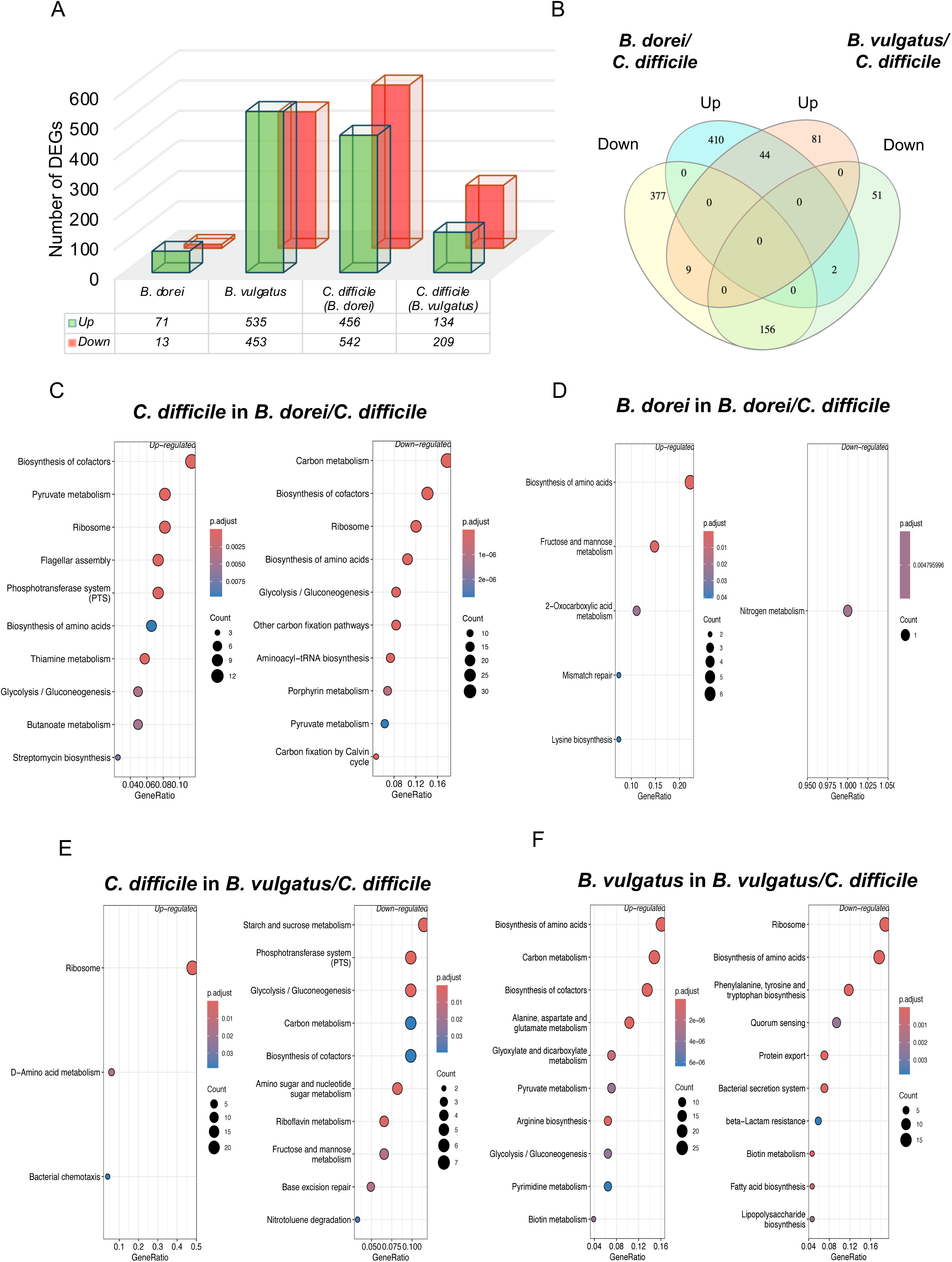
Metabolic shifts occur in *Bacteroides-C. difficile* biofilms. **A**) Numbers of *B. dorei*, *B. vulgatus* and *C. difficile* genes significantly changing in coculture biofilms at 24 h relative to their respective monoculture control biofilms. **B**) Significant differentially expressed *C. difficile* genes that were unique to specific cocultures, or that were overlapping or inversely expressed. Venn diagram was generated using InteractiVenn^102^. KEGG pathway enrichment of significantly expressed genes was carried out using ClusterProfiler, with separate analyses shown for enrichment of *C. difficile* (**C)** or *B. dorei* (**D**) in coculture and for *C. difficile* (**E**) or *B. vulgatus* (**F**) in coculture.

Significant changes were noted in *C. difficile* metabolic pathways which were up- and downregulated in either coculture condition, relative to the monoculture. KEGG analysis revealed distinct shifts in amino acid utilisation in *C. difficile*, showcasing upregulation of pathways involved in biosynthesis of amino acids and cofactors, as well as phosphotransferase systems (PTS) in the presence of *B. dorei*, whilst D-amino acid metabolic pathways were enriched in response to *B. vulgatus* (Fig. 2C,E). *C. difficile* butyrate/butanoate metabolism was also enriched in the presence of *B. dorei,* and closer inspection showed many genes involved in its production were significantly downregulated in the presence of both *Bacteroides* partners (Fig. 2C, Fig. S7A). *C. difficile* PTS pathways exhibited inverse enrichment across cocultures, being upregulated with *B. dorei* but downregulated with *B. vulgatus* (Fig. 2C,E). A higher expression of *C. difficile* PTS genes in the *B. dorei* coculture potentially reflected the metabolic potential of *C. difficile* and the carbohydrate availability in the environment (Fig. S6E-G). Mirroring the increased PTS transcriptional activity, nine CAZyme genes were upregulated in *C. difficile* in the presence of *B. dorei* (Fig. S6H). Substrate prediction using dbCAN3^40^ did not reveal many specific substrates, and only one of these upregulated genes had a predicted lignin substrate (Fig. S6H). PTS gene expression was much lower in the *B. vulgatus* coculture suggesting global metabolic dampening of carbon metabolism with only one fructose PTS gene, *CDR20291_RS11835* and no CAZymes being significantly upregulated (Fig. S6F-H).

Interestingly, genes associated with biofilm formation, such the cell wall proteins *cwp2*, and *cwp66,* were downregulated in the presence of *B. dorei* (Fig. S6D). In contrast, the flagella genes, *flgD* and *fliC,* which have been associated with *C. difficile* biofilm formation^41^, were significantly upregulated in the presence of *B. dorei* (Fig. S6D). Although downregulation of *fliC* was associated with increased biofilm formation of R20291, the role of *fliC* in biofilm formation remains unclear^42,43,44^. Consistent with this, biofilm biomass in *B. dorei/C. difficile* coculture at 24 h was reduced compared to the *C. difficile* monocultures (Fig. S2B). Cocultures with *B. vulgatus* showed relatively higher expression of these genes (Fig. S6D), with a marginal increase in biofilm biomass (Fig. S2B). It was interesting to note an increase in *luxS* (Fig. 3E) which encodes for a quorum sensing regulator, in the presence of *B. dorei.* LuxS mediates production of the quorum sensing molecule autoinducer 2 (AI-2), which was reported to control *B. fragilis*-mediated inhibition of *C. difficile* in dual-species biofilms^39^.

**Figure 3.**
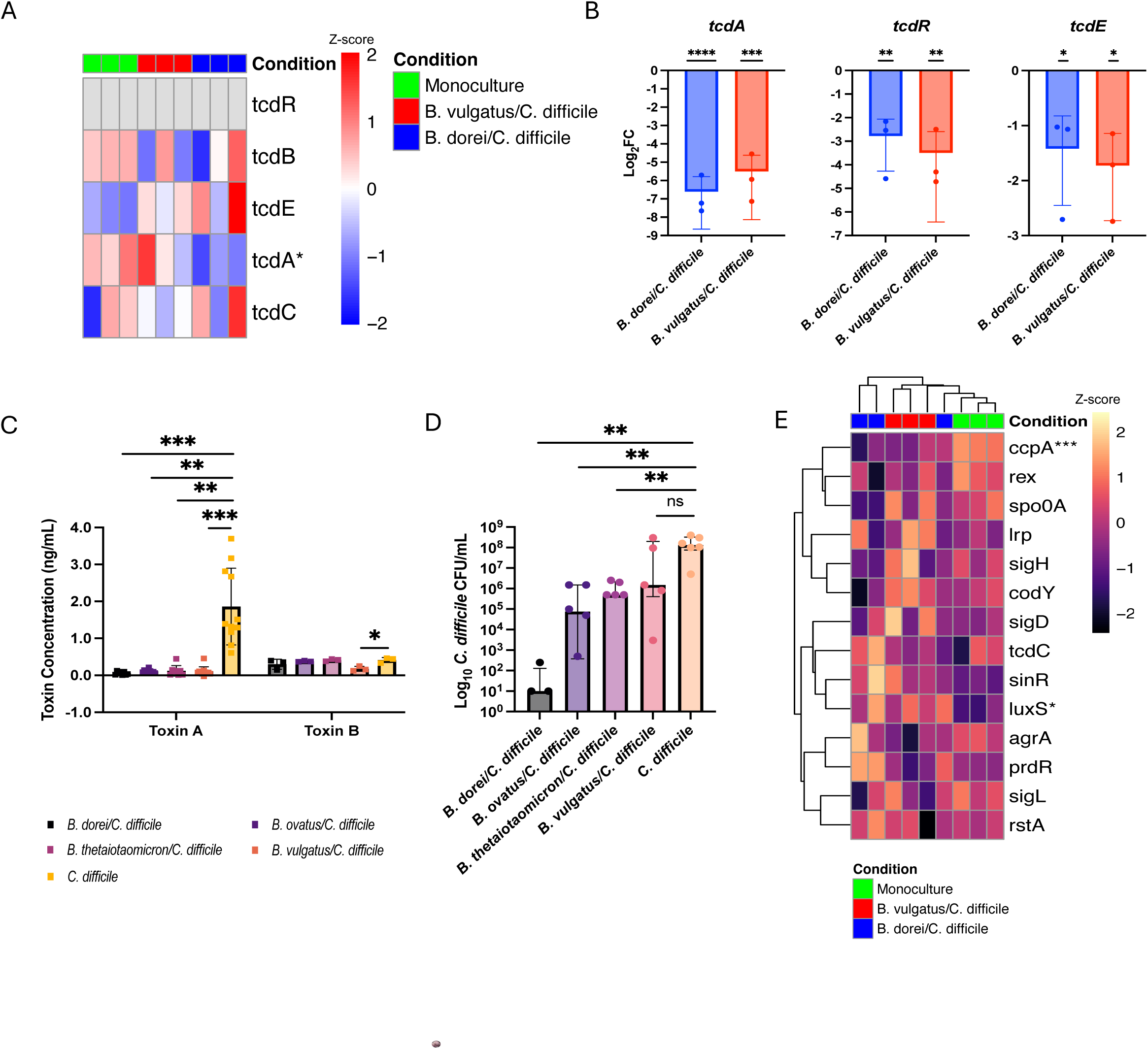
A distinct transcriptional shift in *C. difficile* with notable toxin suppression in *C. difficile*-Bacteroides biofilms. RNA sequencing and downstream analysis were conducted on RNA extracted from dual-species biofilms of *C. difficile* co-cultured with either *B. dorei* or *B. vulgatus* for 24 h. Genes with fewer than 10 counts were excluded from the analysis. **A**) Transcriptional expression of the PaLoc operon in *C. difficile* at 24 h in mono or coculture conditions, presented as a Z-score. **B**) qRT-PCR analysis on coculture and *C. difficile* monoculture biofilms. Log_2_FC of expression of *tcdA*, *tcdE* and *tcdR* genes in the two cultures relative to monoculture. N = 3, mean ± SD. One-sample t-tests were performed from a hypothetical mean of 1. **C**) Toxins from *Bacteroides-C. difficile* coculture biofilm supernatants at 24 h were quantified by ELISA, with a *C. difficile* monoculture biofilm supernatant as a control. Data represents a minimum of 3 biological repeats, mean ± SD is shown. Kruskal-Wallis (toxin A) or one-way ANOVA (toxin B) was carried out, followed by Dunn’s or Dunnett’s multiple comparison test, respectively. **(D)** *C. difficile* CFU/mL from representative coculture and monoculture control biofilms at 24 h. Data represents a minimum of 3 biological repeats, showing the median + interquartile range. Separate Mann-Whitney two-tail tests were performed between coculture and control CFU/mL. **E)** Expression of known regulators of the PaLoc operon in *C. difficile* in different coculture biofilms presented by a Z-score representing high expression (red) and low expression (blue) across samples. For panels **A** and **E**, * denote significant differential expression in *B. dorei/C. difficile* vs *C. difficile* (*) or significance captured in both the *B. dorei/C. difficile* and *B. vulgatus/C. difficile* coculture vs *C. difficile* (***). Significant differentially expressed genes were based on Log_2_FC ≤ -1 and ≥ 1, with padj value < 0.05. Statistical denotations in panels **B-D** represent: ns *P >* 0.05, *\* P <* 0.05*, ** P <* 0.01*, *** P <* 0.001*, **** P <* 0.0001.

Investigation into the transcriptomes of the *Bacteroides* spp. revealed only a nominal number of *B. dorei* genes that were differentially expressed in the presence of *C. difficile* (Fig. 2A), which was likely due to the high variance seen between the replicates (Fig. S6B). On the other hand, *B. vulgatus* displayed large changes in gene expression, with 400 significantly upregulated genes and 500 downregulated genes (Fig. 2A, Fig. S6C). Among the enriched pathways, amino acid biosynthesis was most prominent for both *B. dorei* and *B. vulgatus*, with *B. vulgatus* being specifically enriched for the synthesis of proline, alanine, arginine, and citrulline (Fig. 2D,F; Fig. S.6I). *B. dorei* showed fewer enriched pathways, likely due to the lower number of DEGs. Nevertheless, fructose metabolism (interconversion of D-glucose and D-fructose), and pathways contributing to proline and N-acetyl ornithine production were enriched (Fig. 2D,F; Fig. S6J), mirroring the increased abundance of a *B. dorei* fructose metabolism protein identified by mass spectrometry (Fig. S5B).

Thus, overall, fructose uptake may be central in both cocultures, with an enhanced uptake of fructose by *C. difficile* PTS being potentially a way to scavenge residual nutrients for survival, particularly in a *B. dorei* coculture where *C. difficile* survival is highly impacted.

### *C. difficile* modulates its toxin expression in response *to Bacteroides spp*

Interestingly, the transcriptomic analysis also revealed downregulation of some genes in the *C. difficile* PaLoc operon, which houses genes encoding the primary toxins, TcdA and TcdB (Fig. 3A). This repression was significant for *tcdA* in *C. difficile* cultured with *B. dorei* (Fig. 3A). qRT-PCR analysis confirmed a significant downregulation of *tcdA* when *C. difficile* was cultured with *B. dorei* or with *B. vulgatus* (Fig. 3B). qRT-PCR on other PaLoc genes including *tcdE* and the primary PaLoc regulator, *tcdR* (whose expression were not well captured in the RNAseq due to low reads), also showed significant reduction in coculture with either *Bacteroides* species (Fig. 3B).

To examine if toxin production was impacted, culture supernatants were obtained from 24 h coculture biofilms with *B. dorei, B. vulgatus,* as well as other *Bacteroides* species. All the cocultures showed a potent inhibition of toxin A compared to monoculture biofilms (Fig. 3C). The decrease in toxin levels observed was not dependent on the level of *C. difficile* inhibition, as measured by bacterial counts from representative biofilms (Fig. 3D). No significant changes were observed in toxin levels from planktonic cocultures, but as levels were below the detection range, this was inconclusive (Fig. S8).

Although significant downregulation of *tcdR*, the primary activator of the PaLoc operon, was evident (Fig. 3B), it was pertinent to investigate other regulators including global nutrient regulators, CcpA (glucose), CodY (branch chained amino acids and GTP), PrdR (proline), and Rex (NADH/NAD^+^ ratio), which inhibit toxin expression^45^. Most showed minimal or no significant changes (Fig. 3E). Notably, *ccpA* was significantly downregulated in both cocultures, suggesting that carbon catabolite repression is unlikely to underpin the observed toxin suppression (Fig. 3E). The only *C. difficile* regulator whose expression was significantly upregulated in the presence of *B. dorei* was *luxS* (Fig. 3E); AI-2 has been associated with toxin production, although the underlying mechanisms remain unclear^46–48^. It was also interesting to note that in a *B. dorei* coculture*, prdR* expression was increased (Fig. 3E). Thus, we demonstrate a clear decrease in expression of *C. difficile* toxins when cocultured with *Bacteroides* species as biofilms.

### Toxin reduction is not a result of commensal-mediated secretion inhibition

It was recently reported that *B. thetaiotaomicron* can inhibit release of *C. difficile* toxins^49^. To investigate this possibility, supernatants were retrieved, and bacteria within coculture and monoculture biofilms were lysed by freeze-thawing^50^. Interestingly, *C. difficile* monoculture control biofilms possessed much higher levels of intracellular toxin compared to supernatant and there was a trend of increased toxins in the lysates compared to supernatants in the *B. dorei*, *B. ovatus* and *B. vulgatus* cocultures (Fig. 4A). *C. difficile* colony counts were enumerated in parallel, demonstrating, as expected, a reduction when in coculture with the different *Bacteroides* species (Fig. 4B). Toxin levels in biofilm lysates normalised to *C. difficile* numbers from the respective biofilms showed no significant difference between coculture conditions (Fig. 4C). Normalised supernatant toxin levels demonstrated higher toxins in only the *B. dorei/C. difficile* coculture compared to the control, although a large amount of variation was observed (Fig. 4D). Moreover, the overall % secretion, calculated by dividing the normalised supernatant toxin levels by the total of normalised lysate and supernatant toxins, showed no significant difference between coculture or control conditions (Fig. 4E). Thus, no significant changes were seen in toxin A levels between the lysates and supernatants of *C. difficile* and *C. difficile* cultured with *B. dorei, B. vulgatus, B. ovatus*, or *B. thetaiotaomicron* indicating that the decrease in toxins observed was not due to reduced release or secretion of toxins.

**Figure 4.**
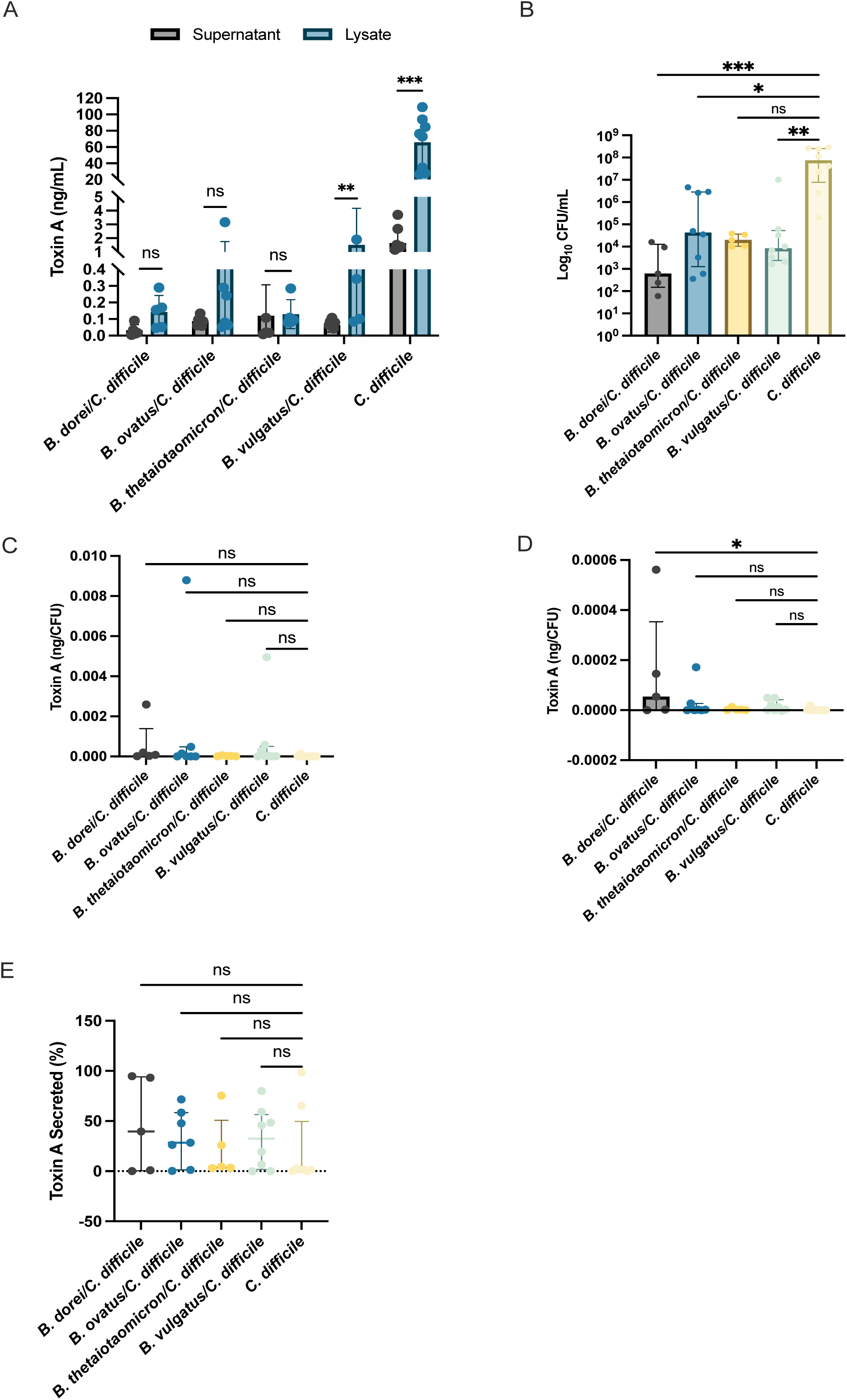
*C. difficile* toxin secretion is not affected by *Bacteroides* in biofilms. **>A**) Quantification of toxin A in supernatants and lysates from monoculture and coculture biofilm cultured for 24 h by ELISA. Data represents the mean ± SD. Individual Wilcoxon matched-pairs signed rank tests/paired t-tests were performed to compare toxin A levels between lysate and supernatant. CFU/mL of *C. difficile* were enumerated in parallel from biofilms used in this experiment **(B**). A Kruskal-Wallis test was performed with Dunn’s correction. Normalised toxin A levels from lysates (**C**) and supernatant samples (**D**), normalised to the CFU of *C. difficile* within each respective biofilm (from B). A Kruskal-Wallis test was performed on (**C**) (*P* = 0.1488) and (**D**) (*P* = 0.0536). A post-hoc Dunn’s test was performed, highlighting a significant difference between the B*. dorei/C. difficile* and the *C. difficile* monoculture control in (**D**) (*P* = 0.0155). E) Toxin secretion was calculated by dividing the normalised supernatant toxin A concentration by the total amount of normalised toxin A in lysate and supernatant together, multiplied by 100. Kruskal-Wallis with Dunn’s correction was performed: no significant difference was observed. * denote statistical significance: ns *P* > 0.05, * *P* < 0.05, ** *P* < 0.01, *** *P* < 0.001. Panel B-E represents the median with the interquartile ranges. All data here represents a minimum of 5 biological repeats.

### Multiomics reveal proline *C. difficile* is cross-fed by specific *Bacteroides* spp. in coculture biofilms

Given the significant changes in metabolism occurring in both the species, we hypothesised that alterations in metabolic pathways may impact toxin production. When we probed the *C. difficile* amino acid pathways, interestingly, we observed significant upregulation of most genes in the *C. difficile prd* operon when cocultured with *B. dorei* (Fig. 5A). A similar trend was also evident in *B. vulgatus* coculture, although the differences were not significant. *prdE*, which shares high structural homology to PrdA^51^, was the only gene in the operon that was significantly upregulated in *C. difficile* cocultured with *B. vulgatus* (Fig. 5A). Analysis of the transcriptome of *B. dorei* and *B. vulgatus*, revealed that the biosynthetic pathway for proline, mediated by ProB, ProA, and ProC^52^, was upregulated in both species (Fig. 5B). These data may suggest that both *Bacteroides* spp. produce proline, and *C. difficile* upregulates its proline reductive pathway in response. The difference in *C. difficile* proline utilisation responses observed by transcriptomics between the two *Bacteroides* species may simply be due to timing of the interaction; *B. dorei* may have already produced proline earlier compared to *B. vulgatus.* Certain Clostridia and *Bacteroides* harbour an alternative to the canonical ProBAC pathway, mediated by the glycyl radical enzyme HypD^53,54^. However, *hypD* expression was not significantly altered in *B. dorei* or *B. vulgatus*, indicating that proline biosynthesis likely occurred primarily through the canonical pathway within coculture biofilms (Fig. 5B). *C. difficile hypD* was significantly downregulated in both *Bacteroides* cocultures, and *proC* also had notably lower expression, in presence of *B. dorei* (Fig. 5A). These findings suggest *C. difficile* may rely more on utilising exogenously derived proline rather than investing in its own biosynthesis.

**Figure 5.**
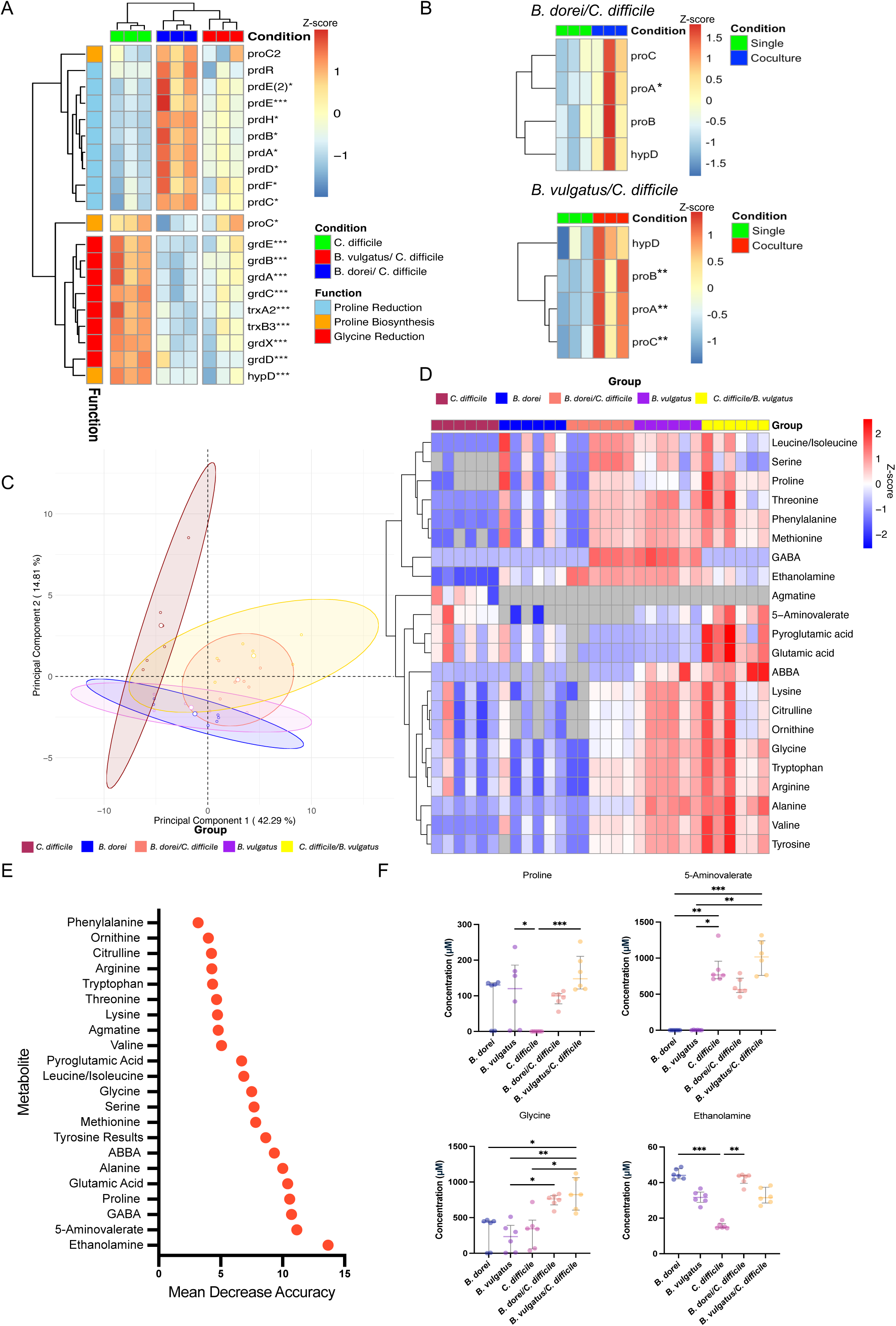
Proline cross-feeding by *Bacteroides* in biofilms. **A**) A heatmap of expression of genes involved in Stickland fermentation pathway in *C. difficile* including proline reductive and biosynthetic pathways enhanced in coculture shown by transcriptomics. B) *B. dorei* and *B. vulgatus* expression of proline biosynthesis. For A) and B), * denote significant differences in the *B. dorei/C. difficile* (*), *B. vulgatus/C. diffiicle* (**) or both (***) cocultures relative to a monoculture. Metabolomics using targeted mass spectrometry was performed on biofilm supernatants from 24 h cocultures and monocultures. **C**) PCA plot of metabolite concentrations between conditions. Concentrations below the limit of quantification (LOQ) were assigned as not applicable (NA). The lowest detected concentration above the LOQ for each amino acid was used for imputation to enable PCA exploration. 95% confidence ellipses are shown. **D**) Heatmap of metabolite concentrations in supernatants of *C. difficile* coculture biofilms with *B. dorei* and *B. vulgatus* and of individual monoculture control biofilms at 24 h. Concentrations were scaled across each row as a Z-score showing higher (red) or lower (blue) concentrations. Amino acid concentrations below LOQ were set as NA, and coloured grey. **E**) Random forest analysis was performed on metabolomics data. Concentrations below the LOQ were imputed based on the minimum concentration found within that amino acid subset that was above the LOQ. Random forest was performed with 500 decision trees, with four splits per tree. **F**) Comparison of the Stickland substrates proline and glycine, Stickland product 5-amino valerate, and amine ethenaloamine, across different conditions, showing median + interquartile range. Concentrations were set to 0 if they fell below the LOQ. A Kruskal-Wallis test was done, and Dunn’s correction was performed to compare between all conditions: *\* P <* 0.05*, ** P <* 0.01*, *** P <* 0.001. Heatmaps in **A**, **B**, and **D** display data presented as a Z score. All data presented here represent N = 6.

To validate this, targeted metabolomics probing a panel of amino acids and amines was performed from *C. difficile-Bacteroides* biofilm supernatants at 24 h. Distinct changes were seen in the metabolite concentrations between conditions, and ∼82% of the variance was captured across the first two main principal components, following imputation (Fig. 5C). Similarly, a separate PERMANOVA analysis revealed that the overall metabolite profiles differed significantly between conditions (F_4,25_ = 20.434, R² = 0.765, *P* > 0.001), and explained ∼76.5% of the variation in the data, with ∼23.5% attributable to residual variation. Distinct clustering patterns of amino acid levels were observed between conditions. Both the *B. vulgatus* monoculture and coculture with *C. difficile* exhibited significantly higher amino acid abundance compared to other conditions (Fig. 5D). The *B. dorei*/*C. difficile* coculture also showed elevated amino acid levels compared to the *C. difficile* monoculture, while the *C. difficile* monoculture generally had lower levels across the panel of metabolites, except for agmatine, which was only present in the *C. difficile* monoculture (Fig. 5D).

To identify the amino acids exhibiting the most substantial changes across conditions, random forest analysis, a form of supervised machine learning, was employed to help identify the amino acids that best discriminate between the five conditions. The topmost discriminating feature, based on the highest mean decrease in accuracy, was the amine ethanolamine which was found in significantly higher abundance in a *B. dorei* monoculture and coculture compared to a *C. difficile* monoculture (Fig. 5E,F). The second highest metabolite was 5-aminovalerate, the end product of proline reduction via Stickland fermentation^55^ (Fig. 5E). Interestingly, proline itself also ranked high among the most discriminating metabolites, ranking fourth (Fig. 5E). Proline levels were notably elevated in the *B. vulgatus*/*C. difficile* coculture compared to the *C. difficile* monoculture, which displayed a marked depletion of proline by 24 h (Fig. 5F). The *B. dorei* monoculture and coculture also showed higher levels of proline, although this difference was not statistically significant (Fig. 5F). High levels of 5-aminovalerate in both cocultures and *C. difficile* monoculture confirms active Stickland fermentation of proline (Fig. 5F). Its notable absence in the *B. dorei* and *B. vulgatus* monoculture also highlights these species do not participate in its fermentation (Fig. 5F). The elevated proline levels in the *B. vulgatus* monoculture and coculture, strongly suggest that *B. vulgatus* supplies proline, likely stimulating the expression of the *prd* operon, suggesting a cross-feeding mechanism via proline. This aligns well with the transcriptomic upregulation of the biosynthetic pathway for proline observed in *B. vulgatus* when in coculture with *C. difficile* (Fig. 5B).

Additionally, glycine concentrations were higher in both the *B. dorei* and *B. vulgatus* coculture compared to all monoculture controls (Fig. 5F). The accumulation of glycine in both cocultures, in contrast to the *C. difficile* monoculture, suggests that *C. difficile* preferentially ferments proline, while glycine fermentation likely remains inactive under these conditions. This is in line with the downregulation of glycine reductase genes observed in *C. difficile* in response to *Bacteroides* partners (Fig. 5A), and fits well with the known inverse relationship between proline and glycine utilisation pathways in *C. difficile*, where the activation of one suppresses the other^56^.

Although proline, glycine and leucine are among the most important Stickland substrates for the reductive pathway^57,58^, certain clostridial species are also capable of utilising a myriad of substrates for Stickland oxidative reactions (leucine, isoleucine, valine, alanine, threonine, tyrosine, methionine) and reductive reactions (proline, glycine, leucine, threonine, phenylalanine) to generate ATP and regenerate NAD+^45,59,60^. Most of these substrates were significantly depleted within a *C. difficile* monoculture biofilm (Fig. S9). In addition to proline and glycine, the Stickland substrates phenylalanine, alanine, tyrosine, valine, threonine and methionine were also found in significantly higher abundance within a *B. vulgatus* coculture (Fig. S9). This suggested that there is an increased pool of amino acids that *C. difficile* may exploit for its survival and continued persistence (Fig. S9).

### *C. difficile* coexists within a microbiota biofilm community with suppressed toxin production

Although we demonstrate crosstalk between specific *Bacteroides* spp. with *C. difficile* via specific metabolic pathways, it is important to understand such interactions in the context of the wider microbiota community. To probe interactions within a human gut microbiota community, we employed a synthetic nine-species biofilm community of representative healthy gut commensals^37^. Individual members of this community can be tracked over time using a qPCR-based approach. We first compared the population dynamics of this commensal biofilm community, which includes three *Bacteroides* species, when cultured in the presence and absence of *C. difficile* over a 48 h period. There were notable changes at the commensal population level in response to *C. difficile* (Fig. S10) although individual commensal species showed similar overall profiles regardless of *C. difficile* status, with *B. adolescentis* and *R. gnavus* staying most abundant (Fig. 6AB). *C. difficile* was inhibited by the microbiota community, with lower *C. difficile* numbers measured across the entire time series compared to *C. difficile* monoculture biofilms (Fig. 6C). However, the overall extent of inhibition remained unchanged over time, with *C. difficile* displaying a reduced yet persistent presence in community biofilms over time (Fig. 6C). To determine whether the *Bacteroides*-associated inhibitory effect and the *Bacteroides* driven *C. difficile* toxin suppression could be recapitulated within this complex commensal community, we performed a multispecies RNAseq analysis from this biofilm community in the presence or absence of *C. difficile*. A complete list of significant differentially expressed genes captured here can be found in Table S3.

**Figure 6.**
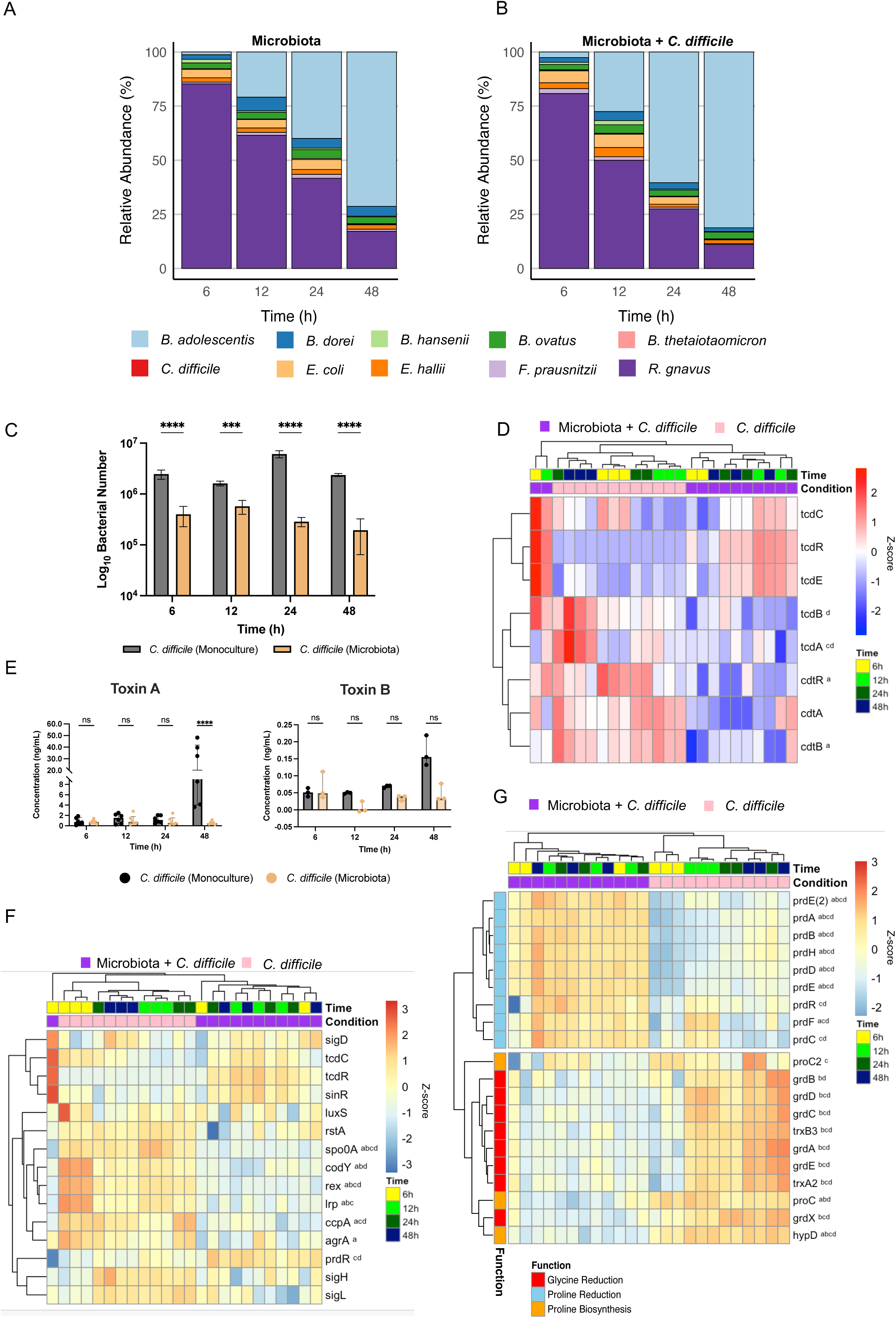
A commensal microbiota community inhibits *C. difficile* proliferation and toxin production through a proline reduction pathway. The relative abundance of each species in the microbiota (**A**) and microbiota + *C. difficile* (**B**) communities, expressed as a percentage across each time point, based on bacterial numbers quantified by PMA-qPCR. N = 3. **C)** Comparison of *C. difficile* number, as predicted by PMA-qPCR, between monoculture and community + *C. difficile* biofilms, over a 48 h time series. N = 3, mean ± SD, A two-way ANOVA was performed, with Šídák’s multiple comparisons correction. **D**) Heatmap of expression of the *C. difficile* PaLoc and CdtLoc operons from the multispecies RNAseq. **E**) Toxins A and B quantified by ELISA over 48 h from biofilms. Data represent a minimum of three biological repeats, displayed as the median + interquartile range. Statistical analysis was performed using multiple Mann-Whitney tests for toxin A and toxin B at each time point. Holm-Šídák correction for multiple comparisons was performed (Toxin A: P = 0.0086). **F**) Expression of known PaLoc regulators over time between *C. difficile* monoculture and *C. difficile* within a commensal community biofilm over time. G) Expression of proline reductase and biosynthetic genes, and glycine reductase genes in C. difficile. For heatmaps in **D**), **F**) and **G**), superscripts denote significantly differentially expressed genes between C. difficile in the microbiota condition relative to the *C. difficile* monoculture control at 6h (^a^),12h (^b^), 24h (^c^) 48h (^d^). Heatmaps in **D**, **F**, **G**, display data as a Z score. * in **C** and **E** denote significance: ns P > 0.05, *** P < 0.001, **** P < 0.0001.

A clear downregulation of *C. difficile* toxin gene *tcdA* was observed at 24 h and 48 h, with significant downregulation of *tcdB* seen at 48 h (Fig. 6D). This was subsequently validated by ELISA quantification (Fig. 6E). Surprisingly, expression of *tcdR and tcdE* appeared to be increased in a community background, although other regulators of the PaLoc like *ccpA*, *codY* and *spo0A* were downregulated (Fig. 6F). Notably, the *prdR*, which is associated with proline metabolism, was significantly upregulated (Fig. 6F), indicating that *prd* activity may be driving the reduction of *C. difficile* toxin production observed here as well. Indeed, genes involved in proline reduction were among the most differentially expressed and consistently upregulated genes over the time course (Fig. 6G). As expected, and mirroring earlier data in the dual-culture RNAseq, the glycine reductase operon was significantly downregulated, as were genes involved in butyrate metabolism, in *C. difficile* cultured in a microbial community, over time (Fig. 6G, Fig. S7B).

It was interesting to note that in addition to the proline pathways, genes involved in a lantibiotic resistance pathway as well as ABC transporter genes associated with antimicrobial resistance were upregulated in *C. difficile* when cultured in the microbiota community. Many top genes upregulated across the time series were identified to be part a putative conjugative transposon (CTn027) that was unique to the 027 strain R20291^61^. This conjugative transposon element was reclassified as Tn6103, an 84.9 kb transposon spans the loci *CDR20291_1740-1809* ^62^. Many genes within Tn6103 region were modulated in the presence of the microbiota community, with significant upregulation of many genes within the Tn6104 region *(CDR20291_RS09510-CDR20291_RS09620 / CDR20291_1744-1764)* which included a putative transcriptional regulator (*CDR20291_RS09525 /CDR20291_1747*), a two-component system (*CDR20291_RS09540-RS09545*/ *CDR20291_1748-1749*), putative lantibiotic ATP Binding Cassette (ABC) transporter system (*CDR20291_RS09550-RS09545/ CDR20291_1751-1752*), a toxin-antitoxin system (*CDR20291_RS09590-RS09595/ CDR20291_1759-1760*), and a MATE-efflux exporter (*CDR20291_RS09760/ CDR20291_1779*).

Taken together our data highlight how a healthy gut microbiota community blocks *C. difficile* proliferation while enabling it to persist in it, shifting its transcriptional priorities toward proline reduction, whilst also upregulating putative antibiotic resistance genes which may contribute toward its survival. Notably, it also underscores the ability of the commensal microbiota to neutralise *C. difficile* toxicity.

## Discussion

The gut microbiota is an important niche where pathogens can co-exist so that they can expand under suitable conditions causing disease. Thus, interactions between a pathogen and individual commensal species likely play a crucial role in preventing active infection. Here, we provide mechanistic insights into how *Bacteroides* species suppress the growth and virulence factor production of the opportunistic human gut pathogen *C. difficile*. We report that the *Bacteroides* species *B. dorei, B. ovatus, B. thetaiotaomicron,* and *B. vulgatus* form dual species biofilms with *C. difficile*, inhibiting *C. difficile* growth within biofilms. Leveraging conventional microbiology techniques alongside multiomics analyses, we demonstrate the inhibitory effects were driven by metabolic shifts, and that proline produced by specific *Bacteroides* species can ‘feed’ *C. difficile*, which in turn blocks the production of toxin A, a key virulence factor that mediates mucosal damage and inflammation. We establish that this nutritional crosstalk occurs within adherent dual-species as well as synthetic multispecies commensal communities.

*Bacteroides* species are highly abundant in healthy human guts. Our findings show that *Bacteroides* species can form biofilms with *B. dorei*, *B. ovatus*, and *B. thetaiotaomicron*, leading to inhibition of *C. difficile* within biofilms, while *B. vulgatus* and *B. fragilis* displayed more variable inhibitory responses. *B. dorei* is one of the most dominant species within the *Bacteroides* genus in the human colon^12^, and may be a key driver of colonisation resistance. We recently reported its inhibitory potential against *C. difficile* in both *in vitro* biofilms and an *in vitro* gut model^37,38^. We also reported a significant *C. difficile* inhibition by a clinical *B. fragilis* isolate in biofilms^39^, although the inhibitory effects were less evident in a biofilm setting for the *B. fragilis* strain 3_1_12 used here. While both strains showed inhibitory effects against *C. difficile* in planktonic culture (Fig. S11A), we observed that the clinical isolate showed significantly faster growth in SAB+ compared to *C. difficile* R20291 (Fig. S11B). This illustrates that strain-specific and medium-specific variations need to be considered when defining such inhibitory effects. The reduced, yet noteworthy impact of *B. vulgatus* on *C. difficile* number is interesting, given its high abundance in the gut and its potential as a candidate for treatment of recurrent CDI due its high engraftment levels^63^.

Cell free supernatants from commensals have been previously reported to have antimicrobial properties^64–67^. In our analysis, a *B. ovatus* monoculture CFS inhibited *C. difficile* (Fig. 1J), mirroring previous findings by Yoon *et al* ^68^. However, due to the absence of bile salts in the present study, the mechanisms of inhibition are likely different. In the case of *B. dorei*, only coculture derived CFS showed inhibitory effects, implying a potential pathogen-triggered product (Fig. 1I). Mass spectrometry, however, primarily identified *C. difficile* proteins in the supernatant (Fig. S5A), suggesting that inhibition may not be due to *B. dorei*-specific secreted proteins, but rather a metabolite or inhibitory small molecule. Further investigations are necessary to reveal any direct inhibitory molecules in these supernatants.

Transcriptomic analysis of pathogen-commensal biofilms indicate that the inhibitory effects observed were driven primarily by distinct metabolic shifts. Distinct patterns were evident for *C. difficile* with the two *Bacteroides* species. A downregulation of *C. difficile* PTS genes was evident with *B. vulgatus*, suggesting it was more reliant on amino metabolism in conditions when inhibitory effects were minimal (Fig. 2E, Fig. S6). However, in a *B. dorei* coculture, where *C. difficile was* highly restricted, induction of *C. difficile* and *B. dorei* genes involved in fructose metabolism (confirmed by mass spectrometry; Fig. S5) and *C. difficile* PTS genes were observed (Fig. 2C, Fig S6E,G). Interestingly, fructose has previously been noted to be a signature of patients with asymptomatic CDI^69^. Thus, under unfavourable growth conditions, *C. difficile* may scavenge for residual extracellular carbohydrates like fructose; fructan may be broken down to fructose by *Bacteroides* which are considered generalist glycan degraders^23^.

Clinical manifestations of CDI are primarily driven by the action of *C. difficile* toxins, toxin A and B, which are important vaccine candidates as well as targets for recent anti-virulence therapeutic strategies^70–73^. A downregulation of toxin gene expression was evident in both *B. dorei* and *B. vulgatus* cocultures, despite their differing effects on *C. difficile* viability (Fig. 3). A previous study showed that *C. difficile* toxin production was reduced when treated with CFS from *B. thetaiotaomicron* monocultures^49^. The study reported that the suppression of toxin release was mediated by the *B*. *thetaiotaomicron*-derived polysaccharide fraction from culture supernatants with no transcriptional impact on toxin expression. The authors claimed that the toxin release was blocked by preventing *C. difficile* autolysis, although the underpinning mechanism was not clear. Additionally, it was unclear if this study factored in the inhibitory effects on *C. difficile* growth^32^. It is possible though that *Bacteroides* cell envelope components may play a role in the CFS effects we report in this study. Direct degradation of *C. difficile* toxins as well as selective killing of *C. difficile* has also been reported by a *Bacillus velezensis* strain ADS024 culture supernatant^49,65,74^. Another study also demonstrated the ability of synthetic community-derived CFS to neutralise toxins by inhibiting host substrate glucosylation^75^. Thus, while we demonstrate that toxin gene repression is mediated through metabolic reprogramming by commensals, other secreted components from commensals may also contribute to supressing *C. difficile* toxins.

*C. difficile* metabolism and toxin expression are known to be tightly linked. The complex regulatory network that controls *C. difficile* toxin expression has environmental cues such as nutrient availability (amino acid and carbon) and redox levels (NAD^+^ levels)^45^. Among many regulators of toxin expression and repression, carbohydrate/sugar availability is a pivotal determinant^76^. In *C. difficile,* PTS sugar availability induces toxin repression via CcpA which binds to the regulatory genes of *tcdA* and *tcdB,* a process known as CCR^76^. Although we observed *C. difficile* modulates expression of specific PTS uptake channels, particularly upregulating fructose channels in the presence of *B. dorei, ccpA* was downregulated in both cocultures (Fig. 3H) thus suggesting CCR was not likely to be involved in toxin repression.

Proline is an essential amino acid required for *C. difficile* growth and has been linked to both toxin levels *in vitro* and implicated in virulence *in vivo*^77–79^. The dual-species transcriptome analysis in this study revealed a clear upregulation of proline biosynthetic pathways in both *Bacteroides* spp., consistent with targeted metabolomics that confirmed increased proline production (Fig. 5), as well as a corresponding increase in the key proline reductase (*prd*) Stickland fermentation pathway, which enables *C. difficile* to utilise proline. Mechanistically, Bouillaut *et al.* proposed that proline abundance activates PrdR, leading to NAD⁺ regeneration and subsequent Rex activation, thereby repressing butyrate metabolism^80^. Increased butyrate levels are known to enhance toxin production^80–82^. The inability of *C. difficile* to ferment proline was shown to reduce toxin levels (*prdB* mutant), yet the incapacity to detect proline levels and regulate the *prd* operon (*prdR* mutant) resulted in increased toxin levels even in high proline environments^56,79^. The expression of *rex* did not change in the coculture biofilms, however it is likely that the activation occurs at a protein level, as suggested previously^80^. Nevertheless, butyrate metabolism was clearly downregulated in *Bacteroides* cocultures, as well as in the synthetic community (Fig. S7).

Bacteroidales strains have been previously shown to accumulate significant amounts of collagen derived proline-based metabolites such as hydroxyproline and proline-hydroxyproline^83^. Studies have also reported amino acid cross feeding between *Bacteroides* and other *Clostridium* species, including proline and formate exchange in *Clostridium ASF356* ^84^. Moreover, carbohydrate limitation has been demonstrated to drive metabolite cross-feeding and coexistence *in vivo* between *B. vulgatus* ATCC 8482 and *C. difficile* R20291, reducing toxin production while enhancing colonisation^85^. In line with our findings, *Bifidobacterium longum* was recently reported to inhibit *C. difficile* growth, as well as toxin production, and using a proteomics approach they demonstrated that proline produced by *B. longum* was utilised by *C. difficile*^86,87^. Thus, proline cross feeding may be a mechanism that multiple commensals in the gut employ to prevent *C. difficile* from producing toxins.

Interspecies competitiveness and cooperativity within synthetic communities have been shown to be important in CDI outcome. In a recent study, a cooperative 4-member consortium protected mice against CDI *in vivo* in part by reducing toxin production, yet had no impact on *C. difficile* burden^87^. Our studies in a complex but defined biofilm community reflected similar mechanisms to that seen in the dual species biofilms, enabling *C. difficile* persistence, yet blocking toxin production. Overall stability of this microbiota community was noted, and although the presence of *C. difficile* had statistically significant impacts on the population dynamics, these were nominal changes (Fig. 6, Fig. S10)^37^. Transcriptomics revealed increased reliance of *C. difficile* on proline fermentation, with downregulation of alternative pathways such as glycine reductase and butyrate synthesis (Fig. 6G, Fig. S7), thus highlighting a central role in Stickland fermentation-mediated toxin repression^87^. While significant reduction of toxin levels was evident, *rex* expression was significantly repressed within the microbiota community biofilm across the entire time series. Thus, toxin downregulation appears to occur independently of *rex* expression suggesting an alternative, yet to be identified, regulatory mechanism in proline-mediated toxin repression, as suggested previously^80^. The importance of proline Stickland fermentation was further demonstrated in a 37-strain consortium, where the inclusion of proline-fermenting strains was required to suppress *C. difficile*^88^. We were unable to demonstrate cross feeding through transcriptomics in the microbiota community. This was mainly due to poor sequencing data from the commensal species within the community and was attributed to technical issues such as high levels of multimapping reads (data not shown). Subsequent analysis using alternative pipelines, such as quasi-mappers like Salmon, that better assign multimapping reads would be useful to probe this further^89,90^. Taken together with our consortium data, this highlights how the tailored design of synthetic commensal consortia can lead to either suppression of *C. difficile* via depletion of essential proline, or by mitigating toxicity via environmental nutritional modulation and induction of proline fermentation.

In summary, we show that in adherent biofilm communities of *Bacteroides* species and *C. difficile*, or among multiple commensal species, *C. difficile* growth and toxin production are inhibited, through modulation of specific bacterial metabolic pathways. Given that proline abundance is positively associated with CDI in humans, and its association with toxin levels both *in vitro* and *in vivo*, our data suggest a commensal-mediated circumvention of *C. difficile* virulence by promoting reliance on proline reduction, resulting in an ablation of toxicity. Such mechanistic understanding of the interactions between individual commensal species is critical, given the growing focus on developing designer microbial cocktails to prevent and treat CDI.

## Materials and Methods

### Bacterial strains and culture conditions

A list of bacterial species and strains can be found in (Table S1). Strains were streaked from 50% glycerol stocks and cultured on Schaedler anaerobe (SAB) broth or agar (Oxoid, UK), routinely supplemented with 2 g/L L-cysteine (Sigma-Aldrich, USA) and 0.005 g/L vitamin K (VWR, USA), which is denoted henceforth as SAB+, and *Clostridium difficile* agar base (Oxoid, UK) (CDA). Strains were routinely cultured under anaerobic conditions at 37 °C in a Don Whitley DG5250 workstation (Don Whitely, United Kingdom).

### Coculture assays and biofilm quantification

Single colonies were inoculated overnight into 10 mL SAB+ broth. After approximately 14-15 h of growth, overnight cultures were normalised to 0.1 OD_600_ in pre-reduced SAB+ liquid media. For cocultures, 0.5 mL of each 0.1 OD_600_ species dilution were mixed, yielding a final concentration of 0.05 OD_600_ for each species within the coculture mix. For single species control biofilms, 0.5 mL 0.1 OD_600_ dilution was mixed with fresh pre-reduced SAB+ media. 1 mL aliquots of each dilution culture were added to pre-reduced 24-well tissue culture treated polystyrene plates (Falcon®, Corning, USA). Wells containing SAB+ alone were used as negative controls. For the multispecies microbiota community, overnight cultures were mixed so that each species had a final concentration of 0.1 OD_600_ within the mixture. For control community cultures without *C. difficile*, fresh SAB+ media was added in its place to ensure no bias. 2 mL aliquots of these cultures were then pipetted into pre-reduced 12-well tissue culture treated polystyrene plates (Falcon®, Corning, USA). At the desired time, biofilms were washed once with PBS. For enumeration of the CFU/mL, the biofilm was subsequently disrupted vigorously in 1 mL pre-reduced PBS, serially diluted, plated onto the appropriate agar plates, and incubated at 37 °C, anaerobically. To help distinguish between single species in coculture, resuspended biofilms were plated onto both SAB+ and CDA plates. For planktonic coculture assays, as with the biofilm assay, overnight cultures were diluted and mixed yielding a final concentration of 0.05 OD_600_ for each species in the mix. 10 mL cultures were set up and incubated for the desired duration. Enumeration of CFU/mL was performed as above.

For biofilm biomass quantification, biofilms were set up in three technical replicates. Wells containing only SAB+ were used as negative controls and blanks for background correction. At the appropriate timepoint, following an initial PBS wash and 30 min drying period, 1 mL filter-sterilised 0.2 % crystal violet (Sigma-Aldrich, USA) was added and incubated for 30 min at 37 °C in anaerobic conditions. Crystal violet was subsequently removed, and the wells were washed thrice with 1 mL PBS, and 1 mL methanol (VWR, USA) was added followed by a 30 min incubation at room temperature (RT). The resulting extracted dye was then diluted 1:1, 1:10 and 1:100 and the OD_570_ was measured in a 96-well plate (CytoOne®Starlab, UK) with a spectrophotometer (SPECTOstar^Nano^, BMG Labtech, Germany).

### Growth curves

Growth curves were set up from overnight cultures, and diluted down to 0.05 OD_600_. Manual growth curves were diluted down into large volumes (i.e. ≥ 40 mL) and 1 ml aliquots were taken at each timepoint. If the OD_600_ exceeded 1, then a subsequent dilution was performed and the OD_600_ remeasured. For automated 24 h growth curves, 300 µL of 0.05 OD_600_ culture was pipetted into pre-reduced 96-well tissue culture treated plates (Starlab, UK) and sealed with ThermalSeal RTS™ sealing films (Sigma-Aldrich, USA). Fresh media was pipetted into empty wells to be used as blanks. The sealed plate was then placed into a Cytation 5 reader (BioTek, Agilent, USA), preincubated at 37 °C and OD_600_ measurements were taken every 15 min, with 10 min shaking and 5 min standing between each read.

### CFS extraction

To obtain CFS, mono- and coculture biofilms were set to a final concentration of 0.05 OD_600_ per species, as previously described. At the 24 h timepoint, supernatants were removed, spun down at 1500 x *g* for 5 min, and filter sterilised through a 0.2 µm filter. CFS were spiked in a 1:1 ratio with overnight *C. difficile* cultures diluted to 0.1 OD_600_ in SAB+. As an internal control for normal growth, fresh SAB+ rather than spent media was used. 1 ml aliquots were pipetted into 24-well pre-reduced plates for biofilms to form for subsequent biofilm assay CFU/mL enumeration and biomass quantification as previously outlined. Growth curves were set up in tandem whereby 300 µL of each final spiked dilution, or control dilution, was pipetted into a pre-reduced 96-well tissue culture treated plate (CytoOne®Starlab, UK), and the OD_600_ was read in a Cytation 5 plate reader (BioTek, Agilent, USA), at 15 min intervals over 24 h, with 10 min shaking and 5 min standing between each read.

### Toxin ELISA

To quantify *C. difficile* toxin A or toxin B within supernatants, an ELISA for the separate detection of each toxin was conducted, following the manufacturer’s instructions (TGC-E002-1, tgcBIOMICS, Germany). For quantification of intracellular toxins within biofilms samples, biofilms were resuspended in 1 mL PBS (containing 0.05% Tween 20 and 1x EDTA-free protease inhibitor (Roche, Sigma-Aldrich, USA). The samples subsequently underwent three consecutive cycles of freeze-thaws, moving the samples from dry ice (-80°C) to 37°C to trigger spontaneous lysis of *C. difficile*^50^. The samples were centrifuged at 2500 x *g* for 5 min, the supernatants moved to new tubes, and these were used for the ELISA, according to the manufacturer’s instructions. Where relevant, toxin levels were also normalised to the CFU/mL of *C. difficile* within the original sample.

### Genomic DNA extraction

Bacterial samples were centrifuged for 5 min, 14,000 RPM, and the subsequent pellets were resuspended in 500 µL Solution A (5 mg/mL lysozyme, VWR, USA), 10 mM Tris, 1mM EDTA, 50mM D-glucose monohydrate (Sigma-Aldrich, USA), and incubated at 37°C for 10 min. Following incubation, 20 µL RNase Solution (20 mg/mL, Invitrogen, UK), Proteinase K (20mg/mL, Invitrogen, UK) and 25 μL 10% sodium dodecyl sulphate (SDS) was added to each sample. Another incubation at 37 °C for 10 min was performed, followed by an additional incubation at 60 °C for 45 min. To each sample, 1 volume phenol-chloroform-isoamyl alcohol (25:21:1) (Sigma-Aldrich, USA) was then added, and briefly vortexed. Samples were subsequently centrifuged at 14,000 RPM for 10 min, and the resulting upper aqueous was removed to a fresh Eppendorf tube. This phenol extraction was repeated and to the resulting removed upper aqueous phase, DNA was precipitated at -20 °C overnight following the addition of 0.1 volume of 3 M sodium acetate, and 1 volume -20 °C 96% ethanol. The samples were then centrifuged at 14,000 RPM for 5 min, the supernatants discarded, and the pellets washed with 200 µL 70% ethanol. The samples were centrifuged again, the ethanol discarded, and the pellets were left to air-dry before resuspending the pellets in an appropriate volume of Tris-EDTA buffer (10mM Tris, 1mM EDTA). PMA-treated samples were routinely resuspended in 40 µL. DNA concentrations were quantified using Qubit dsDNA BR assay kits (Thermo Fisher Scientific, USA).

### PMA-qPCR

Biofilm samples were treated with PMA (Biotium, USA) at a final concentration of 40 µM and incubated in anaerobic conditions in the dark for 10 min. Samples were photoactivated for 15 min by exposing them to blue light (465-575 nm) (PhAST Blue, GenIUL, Spain), prior to DNA extraction. This final concentration of PMA was optimised previously in the Unnikrishnan lab^37^. Primers were designed as previously described^37^ (Table S2). These were designed to amplify the DNA gyrase subunit A (*gyrA*) or topoisomerase I (*topI*) genes of each species. All primers had an annealing temperature between 59-60 °C, with an amplicon size between 100-150 bp. The qPCR was conducted using Luna® Universal qPCR Master Mix (New England Biolabs, USA), on an QuantStudio™3 (Thermo Fisher Scientific, USA), following the manufacturer’s specifications. Plates were sealed with ThermalSeal RTS™ sealing films (Sigma-Aldrich, USA). Melt curves were routinely run at the end of the cycle, following the instrument’s general set-up. Conversion from cycle threshold (C_T_) value to bacterial number was done using standard curves and formulae previously established in the lab^37^.

### RNA isolation and sequence analysis

Dual and microbiota commensal biofilms were cultured in 24- and 12-well tissue culture treated polystyrene plates (Falcon®, Corning, USA), respectively. Biofilms were resuspended in 750 µL LETS buffer (0.1M LiCl, 0.01M Na_2_EDTA, 0.01M Tris-HCl (pH 7.5), 0.2% SDS) for subsequent TRIzol-based extraction. The resuspensions were then homogenised in 2 mL Lysing Matrix B (MP Biomedicals, USA) tubes, by bead beating for 6 cycles at 6.5 m/s for 40 seconds in a FastPrep-24-5G (MP Biomedicals, USA). Samples were kept on ice for 3 min between each cycle. The samples were subsequently spun at 15,000 *x g* for 10 min, and the supernatants were transferred into 2 mL nuclease-free Biosphere screw-cap tubes (Sarstedt Ltd., Germany) prior to extraction.

A 1 mL aliquot of TRIzol Reagent (Invitrogen, USA) was added to each supernatant, followed by a 5 min incubation at RT. 200 µL chloroform (Sigma-Aldrich, USA) was added to each sample and briefly vortexed and incubated again at RT for 5 min. Following this incubation, the samples were centrifuged at 12,000 *x* g at 4°C for 15 min. The resulting upper aqueous phases were then removed to new tubes, 500 µL isopropanol was added to each tube, and the samples were incubated at RT for 10 min followed by a 10 min centrifugation at 12,000 *x* g at 4 °C. The supernatants were discarded, and the pellets were allowed to air-dry prior to resuspending them in 30 µL RNase-free water. RNA concentrations were then quantified using Qubit RNA BR assay kits (Thermo Fisher Scientific, USA). RNA samples were DNase treated using the Turbo DNA-free kit (Thermo Fisher Scientific, USA), following the rigorous treatment according to the manufacture’s protocol. Briefly, samples were, where necessary, diluted to a final concertation of 200 ng/µL in a total reaction volume of 50 µL. Each 50 µL reaction contained 5 µL 10x Turbo DNase Buffer and 1 µL DNase. Upon addition of DNase, samples were gently mixed, and incubated at 37 °C for 30 min. 10 µL DNase Inactivation Reagent was added to each tube, gently mixed by flicking and incubated at RT for a further 5 min. The tubes were flicked at 2 min intervals to ensure the inactivation reagents remained well dispersed in solution. Finally, the samples were centrifuged for 1.5 min at 10,000 x *g*. The supernatants were then moved to new nuclease-free tubes. To further remove contaminants such as residual DNA, proteins or carbohydrates, samples were treated with lithium chloride. Here, LiCl Precipitation Solution (Invitrogen, USA) was added to each DNase-treated RNA sample at a final concentration of 2.5 M. The samples were then incubated at –20 °C for 30 min, prior to centrifugation at 14,000 RPM at 4 °C for 15 min. The supernatants were discarded, the pellets washed with 200 µL ice-cold 70 % ethanol and were then subsequently centrifuged for 5 min at 14,000 RPM, before air-dying the pellets and resuspending them in 20 µL nuclease-free water. The samples were then quantified for both DNA and RNA using Qubit BR DNA and RNA kits (Thermo Fisher Scientific, USA).

Total RNA was then sequenced by GENEWIZ (Azenta Life Sciences, USA) who performed rRNA depletion, cDNA synthesis, library preparation, and subsequent paired-end sequencing. The minimum read depth for mono- and cocultures were set at 10 million. Multispecies microbiota community samples were sequenced at a minimum depth of 40 million. Read lengths were 150 bp PE-sequencing. The resulting FastQ file quality was assessed using FastQC (Version. 0.12.1)^91^. Reads were trimmed to remove adapter contaminants using Trimmomatic (Version 0.39)^92^, based on a list of common adapter sequences. Unpaired reads were ignored. FastQC was rerun to ensure satisfactory quality of output files. Following trimming, reads were aligned to the appropriate genomes using HISAT-2 (Version 2.1.0)^93^ : single genomes for single-species samples, or concatenated genomes of the relevant species for coculture or community samples. This was done by concatenating the genome files in Linux. Accession numbers for the respective genomes can be found in Table S4. Prior to running HISAT-2, single and concatenated genomes were indexed using HISAT-2 Build. As reads came from strictly bacterial cultures, the in-built splice-awareness setting was turned off in HISAT-2. Unpaired reads were ignored during this step. The resulting Sequence Alignment Map (SAM) files were converted to Binary Alignment Map (BAM) files, subsequently sorted, and indexed, using Samtools View, Samtools Sort, and Samtools Index commands, respectively^92^. Unmapped reads were discarded during this step. Finally, to get gene counts from the reads, LiBiNorm count (Version 2.4)^94^ was performed in HTSeq-count-compatible mode, using the relevant genome annotations. Again, for coculture or community samples, concatenated annotations were generated and used. For multispecies samples, the count files were demultiplexed to extract species-specific counts from each sample, enabling downstream analysis using the R package DESeq2 (Version 1.40.2)^95^. Significant differential gene expression were set at the threshold of *P* < 0.05, Log_2_FC of ≤ -1 and ≥ 1. Genes with fewer than 10 counts were excluded from the analysis.

### KEGG Enrichment analysis

Kyoto Encyclopaedia of Genes and Genomes (KEGG) pathway analysis was conducted using clusterProfiler (Version 4.8.3)^96^ package in R. Due to the poor annotation of KEGG numbers within a lot of the annotation files across the species used, InterProScan (Version 5.59-91.0)^97^ was run on the protein fasta files of each genome to assign GO terms, whilst GhostKoala (Version 2.0)^98^ was run on each genome protein fasta file to generate a more robust list of KEGG numbers. Protein files were retrieved from using the same accession numbers for each species as listed in Table S4. The relevant KEGG numbers were assigned to the appropriate gene IDs by mapping gene ID to protein ID in the Gene Transfer Format (GTF) genome annotation file, and then each gene ID was linked to its corresponding KEGG numbers using the protein ID-to-KEGG associations provided by the GhostKOALA and InterProScan outputs. KEGG maps were generated using KEGG Mapper^99^.

### Carbohydrate-active enzyme gene annotation

Genome protein fasta files were annotated for carbohydrate-active enzymes (CAZymes) using HMMER: dbCAN3-sub tool (e-value < 1e-15; coverage > 0.35) in the dbCAN3 meta server^40^. dbCan3 was also used to predict predicted glycan substrates of these proteins^100^. CAZyme-associated proteins were then mapped to their associated gene ID to explore their transcriptional activity.

### RT-qPCR

From DNase-treated RNA samples, cDNA was generated using SuperScript IV Reverse Transcriptase (Thermo Fisher Scientific, USA), according to manufacturer’s instructions. RT-qPCR was conducted using Luna® Universal qPCR Master Mix (New England Biolabs, USA). Expression of target genes using primers listed in Table S2 was normalised to the housekeeping *C. difficile gyrA* gene using the ΔΔCt method.

### Metabolomics

Supernatants were removed from 24 h biofilms, centrifuged at 15,000 x g for 5 min and sterilised through a 0.2 µM, prior to storing at -80 °C and sending them for mass spectrometry. Supernatants were diluted 1:10 in water and subsequently extracted by protein crash with the addition 120 µl of extraction buffer (a 1:1 mixture of acetonitrile (VWR 83640.320) and methanol (VWR 83638.320) containing 0.1% formic acid (Fisher A117-50) and 20 µM of each of the following: amoxicillin (Sigma-Aldrich/Merck A8523), caffeine (Fluka A8523), ibuprofen and donepezil as internal standards), mixing briefly by shaking (15 s, 1500 RPM), incubating at 4°C for approximately 30 min and centrifuging (5 min, 3200 *x* g, 4°C). 15 µl of cleared extract was pipetted off into two 384-well PCR plates for quantification of amino acids and amines using targeted liquid chromatography coupled mass spectrometry (LC-MS). All LC-MS measurements were carried out on an Agilent Infinity 1290 LC coupled to an Agilent 28 6470B triple quadrupole mass spectrometer with JetStream ion source. Multiple reaction monitoring (MRM) was used to monitor compound-specific precursor-fragment transitions (usually two per compound). A dilution series of a mixed analytical standards, water blanks and a QC sample (created by pooling all the samples) were injected in regular intervals. Amino acids were measured using a using a low-pH hydrophilic interaction chromatography (HILIC) method with a ACQUITY UPLC BEH Amide 1.7um 2.1 x 100mm column (Waters 186004801), as described previously [https://www.sciencedirect.com/science/article/pii/S0092867416312375?via%3Dihub, https://cshprotocols.cshlp.org/content/2017/9/pdb.prot089094]. Amines were measured using a shortened low-pH HILIC method adapted from https://www.sciencedirect.com/science/article/pii/S0039914020311590?via%3Dihub using an ACQUITY BEH Amide 1.7uM, 2.1 x 50 mm (Waters 186004800) column.

Raw data were analysed using MassHunter Quantitative Analysis v10.1. Compounds which were absent in all samples, had unacceptable peak shapes, or were detected at concentrations lower than the lowest calibration standard in all samples, were excluded from further analysis. Metabolite concentrations were calculated based on 9 point calibration curves using blank offset. All raw data, LC-MS acquisition methods and MassHunter quantification methods have been deposited in a Mendeley data repository (DOI: 10.17632/cvt5kmp3mb.1) and will be made publicly available upon publication.

Values below the limit of quantification (LOQ) were assigned as NA. For the PCA exploratory analysis and Random Forest (RF) analysis, NA values were imputed using the minimum concentration identified for each amino acid across the database. In the overall statistical comparison of concentrations, values falling below the LOQ were assigned an arbitrary value of 0. To evaluate the importance of each metabolite in distinguishing between all the conditions, RF analysis was performed. In this analysis, imputed amino acid concentrations were used where necessary, and supervised machine learning was applied using the RF package in R (Version 4.7.12)^101^, with 500 trees, and 4 splits per tree.

### Statistical Methods

Statistical analysis was performed in either R v2023.06.0 or Graphpad v10.3.1. perMANOVA analysis was accomplished using the vegan R package^102^. All data were subjected to a Shapiro-Wilko normality test to assess data distribution. For parametric analyses unpaired/paired T-tests (pairwise comparisons), one-way or two-way ANOVAs (grouped dataset comparisons), and one-sample T tests (comparison against a reference value e.g. fold change data) were employed. For nonparametric analyses Mann-Whitney U/ Wilcoxon matched-pairs signed rank tests (pairwise comparison), Kruskal-Wallis tests (grouped dataset comparisons), and Wilcoxon signed-rank tests (single-sample comparison against a reference value) were used. Where applicable, post hoc tests were performed to correct for multiple comparisons. Complete description of analyses used is outlined in each figure legend.

## Funding

We would like to thank the Wellcome trust (grant number 227934/Z/23/Z), the BBSRC International Partnership fund, and the Warwick A*STAR research attachment programme for providing funding for this work. KRP and SKA acknowledge funding from European Research Council (ERC) under the European Union’s Horizon 2020 research and innovation programme (grant no. 866028).

## Acknowledgements

We acknowledge Dr. Cleidiane Zampronio (WPH Proteomics Facility RTP, University of Warwick) for performing the proteomics mass spectrometry and thank Stephan Kamrad, University of Cambridge for input on metabolomics measurements

## Author Information

K.K.B.B.: Conceptualisation, methodology, investigation, formal analysis, and writing (original draft and editing).

M.U.: Conceptualisation, methodology, supervision, formal analysis, funding and writing (review and editing).

S.K.A.: Metabolomics measurements and raw data processing.

N.N.: Conceptualisation, supervision, funding and writing (review).

K.R.P.: Investigation, supervision, and writing (review).

## Competing interests

The authors declare no competing interests.

---

**Figure S1. Growth kinetics of species in SAB+ or in presence of cell free supernatant (CFS). A**) Growth curves of various species in SAB+ over a 12 h period based on OD_600_. Using the growth curve data, growth rates, N= 3, mean ± SD (**B)** and doubling times (**C**) were calculated using the R package Growthcurver^103^. A one-way ANOVA was performed on **B** which followed by Holm-Šídák correction, mean ± SD shown. A Kruskal-Wallis test was performed on **C**, followed by Dunn’s correction, median with the interquartile range is shown. **D**) Inoculums of species used biofilms were enumerated from cocultures of *Bacteroides spp*. and *C. difficile*. A Kruskal-Wallis test was performed with Dunn’s correction showing no significant different in bacterial numbers. Median with the interquartile shown. **E**) Growth curves of *C. difficile* in SAB+ spiked with 50% CFS derived from different *Bacteroides* monoculture biofilm cultured for 24 h. N = 3, mean ± SEM is shown. ns *P* > 0.05.

**Figure S2. Commensal association can affect *C. difficile* biofilm formation**. Monoculture and coculture biofilms were stained with 0.2% crystal violet at 12 h, 24 h and 48 h. **A**) Representative images of some mono and coculture biofilms formed at 12 h, stained with 0.2 % crystal violet. **B**) Biofilm biomass was compared across different times; blank corrected OD_570_ was compared between conditions. Data represents a minimum of 3 biological replicates, mean ± SD shown. Two-way ANOVAs were performed followed by Tukey’s multiple comparisons test. **C**) Fold change in biomass (OD_570_) of each coculture at 12 h and 24 h, standardised to the biomass (OD_570_) of a *C. difficile* monoculture. N = 3, mean ± SD shown A one-sample t-test was performed on each, from a hypothetical mean of 1. **D)** Biofilm biomass (OD_570_) from *C. difficile* spiked with coculture CFS at 24 h was compared to one treated with a control *C. difficile* monoculture biofilm CFS. N = 4. A one-way ANOVA with Dunnett’s correction was performed. Statistical significance is denoted by: ns *P* > 0.05, * *P* < 0.05, ** *P* < 0.01, *** *P* < 0.001, **** *P* < 0.0001. Each biological replicate was the average of three technical replicates.

**Figure S3. *Bacteroides*-mediated inhibition is *C. difficile* strain-specific** A) Colony counts of *Bacteroides* spp. and *C. difficile* 630 from mono- and coculture at 24 h. N = 4, mean ± SD shown. Two-tailed Mann-Whitney or unpaired t-tests were performed to compare each respective monoculture vs coculture counts. **B**) Growth curves of single species as measured by OD_600_ over 12 h. N = 3, mean ± SD is shown. **C**) Growth rates were calculated from growth curve data generated in (**B**) using Growthcurver. Median + interquartile range is shown. A Kruskal-Wallis test was performed, followed by Dunn’s correction. ns *P* > 0.05, * *P* < 0.05.

**Figure S4. *Bacteroides*-mediated inhibition of *C. difficile* is not as potent in planktonic milieu. A**) Bacterial numbers in the inoculum was measured by CFU/mL, from *Bacteroides spp./C. difficile* cocultures and respective single species monocultures. N = 3, mean ± SD is shown. A one-way ANOVA or Kruskal-Wallis test was carried out with Tukey’s or Dunn’s correction, respectively. **B)** Monoculture and cocultures from A) were grown planktonically over 48 h, and CFU/mL were enumerated. N = 3, median + interquartile range is shown. Where bacterial numbers were below the detection limit, an arbitrary value of 10 was imputed to allow for statistical analysis. Multiple Mann-Whitney tests were performed, comparing each species CFU/mL between coculture and monoculture, at each respective timepoint. ns *P* > 0.05.

**Figure S5. Mass spectrometry from *B. dorei/C. difficile* biofilm coculture and monoculture biofilm supernatants at 24 h. A**) Significantly differentially abundant proteins identified by mass-spectrometry from supernatants of *B. dorei*-*C. difficile* coculture biofilms after 24 h, relative to monoculture biofilm supernatants. A two-sample t-test was performed. False Discovery Rate (FDR) was set to 0.05 and s0 was set as 0.1 (default for Perseus). Significant genes that passed these cut-offs are coloured in red, with non-significant proteins coloured in blue. **B-C**) KEGG pathways for PTS pathways enriched in *C. difficile* (**B**) and fructose and mannose metabolism enriched in *B. dorei* in, coculture (**C**). DEPs that were significantly more abundant in the *B. dorei/C. difficile* coculture are coloured in green, those that were significantly more abundant in the monocultures are coloured in red. N = 3.

**Figure S6. Heatmaps and pathway maps showing expression profiles from 24 h biofilms.** A-C) Heatmaps showing overall expression profiles from 24 h biofilms from RNAseq data presented as a Z-score for *C. difficile* in different coculture conditions and monoculture control (**A**), and *B. dorei* expression (**B**) or *B. vulgatus* expression (**C**) when cultured on its own or with *C. difficile*. **D**) Expression of *C. difficile* genes associated with biofilm formation within different monoculture and coculture biofilms at 24 h. E-F) KEGG maps of significantly differentially expressed *C. difficile* PTS pathways in *B. dorei* (**E**) and *B. vulgatus* (**F**) coculture biofilms. **G**) Heatmap (presented as a Z-score) showing significant differential expression of *C. difficile* PTS genes in different coculture biofilms, annotated with the respective transported carbohydrate. **H**) Heatmap (presented as a Z-score) of *C. difficile* CAZyme genes differentially expressed in different coculture biofilms, along with their predicted glycan substrates, as predicted by dbCan3**. I-J)** KEGG maps of *B. vulgatus* (**I**) or *B. dorei* (**J**) genes involved in amino acid biosynthetic pathways that are significantly differentially expressed in coculture with *C. difficile* in biofilms. E,F,I,J: significantly upregulated or downregulated genes were coloured in green and red, respectively. For heatmaps A-D,G and H, * denote significant differential expression in *C. difficile* when cocultured with *B. dorei* (*) or *B. vulgatus* (**) relative to the monoculture. *** denotes significant differential expression in both cocultures relative the monoculture condition. Log_2_FC ≤ -1 or ≥ 1, padj < 0.05.

**Fig. S7. *C. difficile* butyrate metabolism is downregulated in a *Bacteroides* and microbiota community cocultures.** Graphs display the Log_2_FC from RNAseq of some *C. difficile* genes involved in butyrate metabolism in either the *Bacteroides* coculture relative to a *C. difficile* monoculture biofilm (**A**), or a commensal community + *C. difficile* biofilm, relative to a control commensal community biofilm at each respective timepoint (**B**). * denotes significant differential expression (Log_2_FC ≤ -1 or ≥ 1, padj < 0.05).

**Figure S8. Toxin levels do not differ in planktonic cocultures.** Toxin concentrations quantified from planktonic cocultures and *C. difficile* monocultures at 24 h. A Kruskal-Wallis or one-way ANOVA were performed on toxin A (*P* = 0.2549) and toxin B (*P* = 1.431), respectively. N = 3, mean ± SD shown. All samples fell below the detectability cut-off. ns *P* > 0.05.

**Figure S9. Concentrations of Stickland substrates as quantified by mass spectrometry.** Metabolomic analysis of supernatants from various monoculture and coculture biofilms at 24 h. For concentrations that were NA, an arbitrary value of 0 was assigned to allow for statistical analyses. A Kruskal-Wallis or One-way ANOVA analysis was performed to compare between conditions, followed Dunn’s or Tukey’s multiple comparison correction, respectively. N = 6. All graphs display the median + interquartile range. * *P* < 0.05, ** *P* < 0.01, *** *P* < 0.001, **** *P <* 0.0001.

**Figure S10. Bacterial numbers of individual species within the commensal community *±* C. difficile** Bacterial numbers within multispecies biofilms cultured in presence or absence of *C. difficile* were calculated using PMA-qPCR at 6-48 h. A-B represent the calculated number of bacteria per species over time in the microbiota (**A**) and microbiota + *C. difficile* community (**B**) as deduced by PMA-qPCR. A two-way ANOVA was performed on each, followed by a Tukey’s multiple comparison correction. The a and b denotations represent significance which can be found in panel C. **D**) Bacterial numbers of individual species within the commensal community ± *C. difficile* over time were compared as deduced by PMA-qPCR. Two-way ANOVAs were performed. Šídák correction was performed for comparisons between condition at each respective timepoint. All data presented is of three biological replicates, each with three technical replicates, mean ± SD is shown. ns *P* > 0.05, * *P* < 0.05, ** *P* < 0.01, *** *P* < 0.001, **** *P* < 0.0001.

**Figure S11. Comparison of two strains of *B. fragilis* with *C. difficile* in biofilm and planktonic coculture. A)** *B. fragilis* strain 3_1_12 or a clinical isolate^39^ were cultured with *C. difficile* in biofilms or planktonic culture over 48 h, as measured by CFU/ml. Data represents a minimum of three biological repeats. Median plus interquartile range is shown. Multiple unpaired t-tests showed no significant differences. **B**) Planktonic growth curves over 12 h were performed and growth rates were calculated using Growthcurver. N = 3, mean ± SD is shown. A one-way ANOVA followed by Dunnett’s multiple comparison test was performed. ns *P* < 0.05, * *P* < 0.05.

**Table S1. List of main bacterial species and strains used.**

**Table S2. List of primers used in this study for RT-qCPR and PMA-qPCR.**

**Table S3. List of significantly differentially expressed genes from the coculture and multispecies RNAseq experiments.**

**Table S4. Reference genomes and annotations used in analyses.**

## Notes

### Competing Interest Statement

The authors have declared no competing interest.

