## Supplemental figures S1-S11 for "*Bacteroides*-driven metabolic remodelling suppresses *Clostridioides difficile* toxin expression in mixed biofilm communities"

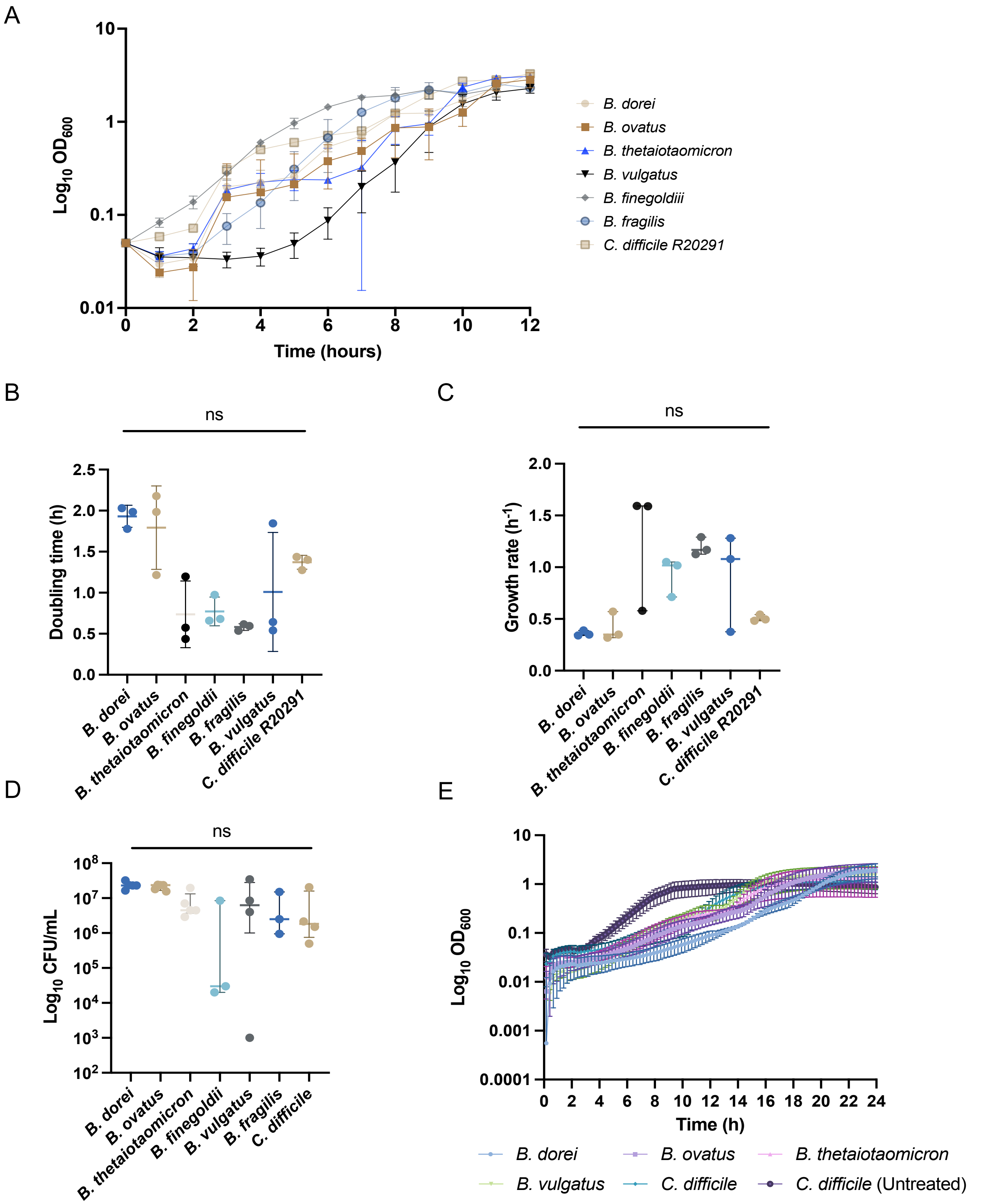


Figure S1. **Growth kinetics of species in SAB+ or in presence of cell free supernatant (CFS)**. **A**) Growth curves of various species in SAB+ over a 12 h period based on $OD_{600}$. Using the growth curve data, growth rates, N= 3, mean ± SD (**B**) and doubling times (**C**) were calculated using the R package Growthcurver[103]. A one-way ANOVA was performed on **B** which followed by Holm-Šídák correction, mean ± SD shown. A Kruskal-Wallis test was performed on **C**, followed by Dunn's correction, median with the interquartile range is shown. **D**) Inoculums of species used biofilms were enumerated from cocultures of *Bacteroides spp*. and *C. difficile*. A Kruskal-Wallis test was performed with Dunn's correction showing no significant different in bacterial numbers. Median with the interquartile shown. **E**) Growth curves of *C. difficile* in SAB+ spiked with 50% CFS derived from different *Bacteroides* monoculture biofilm cultured for 24 h. N = 3, mean ± SEM is shown. ns *P* > 0.05.

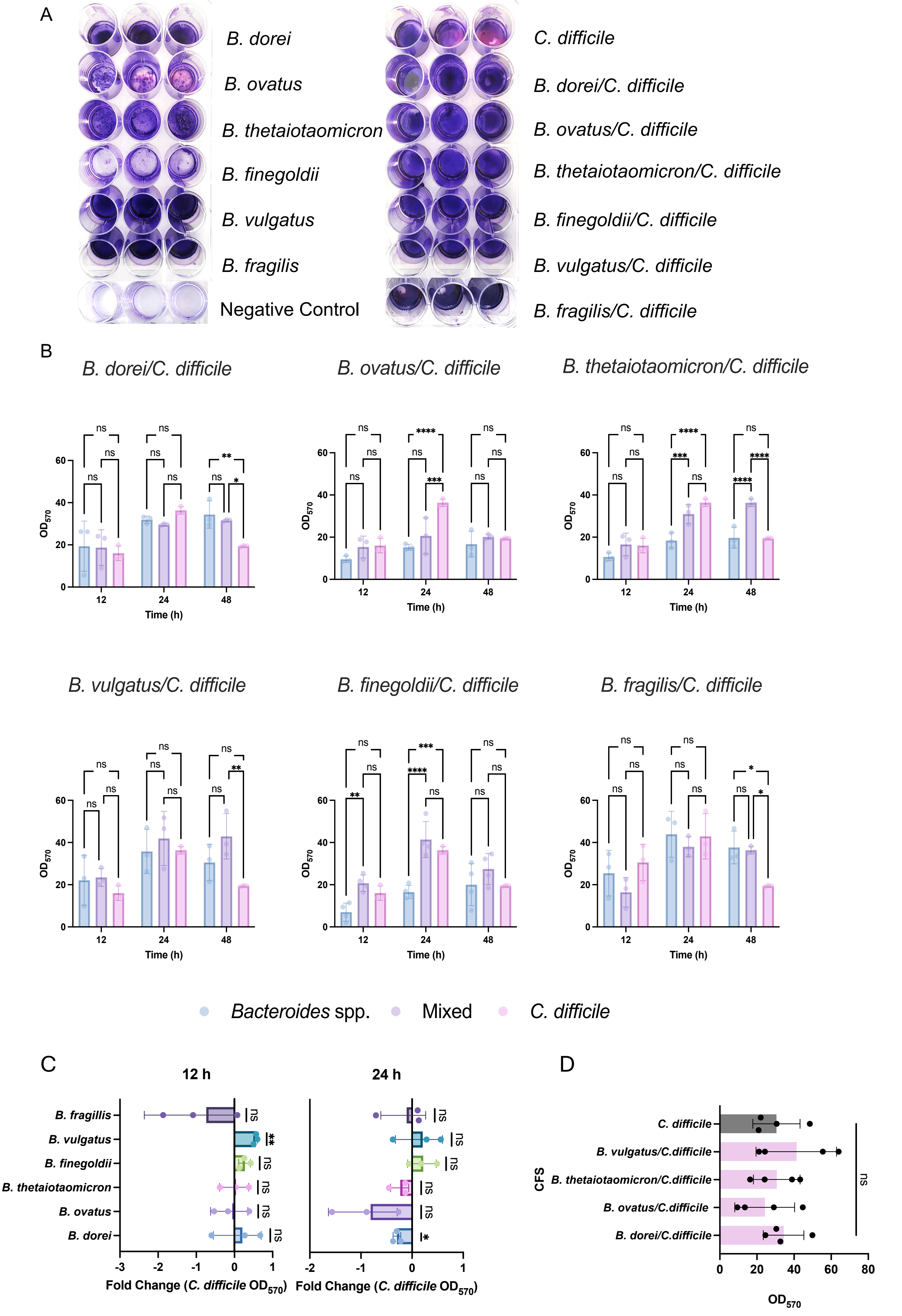

A
B. dorei
B. ovatus
B. thetaiotaomicron
B. finegoldii
B. vulgatus
B. fragilis
Negative Control
C. difficile
B. dorei/C. difficile
B. ovatus/C. difficile
B. thetaiotaomicron/C. difficile
B. finegoldii/C. difficile
B. vulgatus/C. difficile
B. fragilis/C. difficile
B
B. dorei/C. difficile
B. ovatus/C. difficile
B. thetaiotaomicron/C. difficile
B. vulgatus/C. difficile
B. finegoldii/C. difficile
B. fragilis/C. difficile
OD570
Time (h)
12
24
48
Bacteroides spp.
Mixed
C. difficile
C
12 h
24 h
B. fragillis
B. vulgatus
B. finegoldii
B. thetaiotaomicron
B. ovatus
B. dorei
Fold Change (C. difficile OD570)
D
C. difficile
B. vulgatus/C.difficile
B. thetaiotaomicron/C.difficile
B. ovatus/C.difficile
B. dorei/C.difficile
CFS
ns
OD570

Figure S2. **Commensal association can affect *C. difficile* biofilm formation**.
Monoculture and coculture biofilms were stained with 0.2% crystal violet at 12 h, 24 h and 48 h. **A**) Representative images of some mono and coculture biofilms formed at 12 h, stained with 0.2 % crystal violet. **B**) Biofilm biomass was compared across different times; blank corrected $OD_{570}$ was compared between conditions. Data represents a minimum of 3 biological replicates, mean ± SD shown. Two-way ANOVAs were performed followed by Tukey's multiple comparisons test. **C**) Fold change in biomass ($OD_{570}$) of each coculture at 12 h and 24 h, standardised to the biomass ($OD_{570}$) of a *C. difficile* monoculture. N = 3, mean ± SD shown A one-sample t-test was performed on each, from a hypothetical mean of 1. **D)** Biofilm biomass ($OD_{570}$) from *C. difficile* spiked with coculture CFS at 24 h was compared to one treated with a control *C. difficile* monoculture biofilm CFS. N = 4. A one-way ANOVA with Dunnett's correction was performed. Statistical significance is denoted by: ns $P > 0.05$, * $P < 0.05$, ** $P < 0.01$, *** $P < 0.001$, **** $P < 0.0001$. Each biological replicate was the average of three technical replicates.

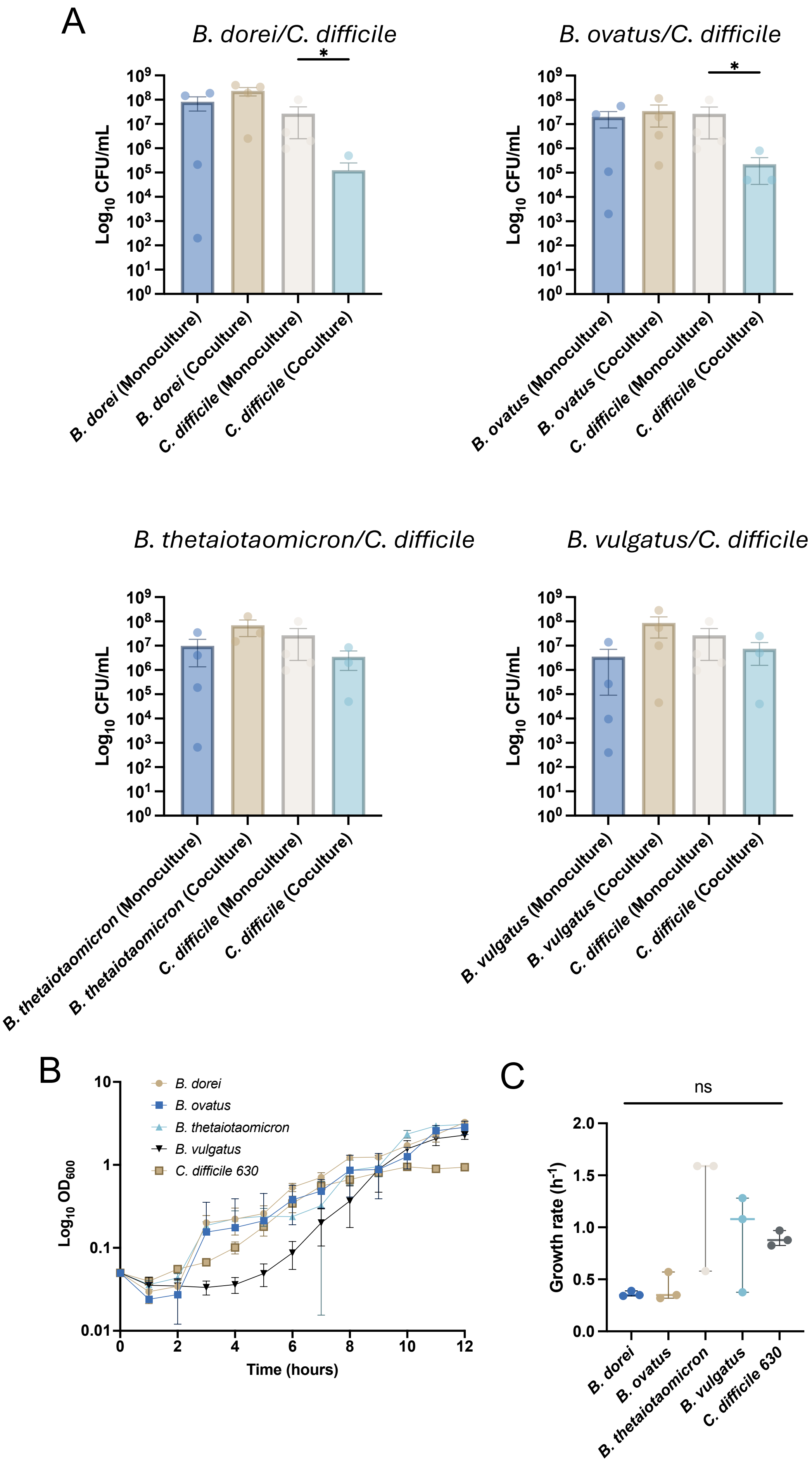
A
B. dorei/C. difficile
B. ovatus/C. difficile
B. thetaiotaomicron/C. difficile
B. vulgatus/C. difficile
*
Log10 CFU/mL
B. dorei (Monoculture)
B. dorei (Coculture)
C. difficile (Monoculture)
C. difficile (Coculture)
B. ovatus (Monoculture)
B. ovatus (Coculture)
B. thetaiotaomicron (Monoculture)
B. thetaiotaomicron (Coculture)
B. vulgatus (Monoculture)
B. vulgatus (Coculture)
B
B. dorei
B. ovatus
B. thetaiotaomicron
B. vulgatus
C. difficile 630
Log10 OD600
Time (hours)
C
ns
Growth rate (h-1)
B. dorei
B. ovatus
B. thetaiotaomicron
B. vulgatus
C. difficile 630

Figure S3. ***Bacteroides*-mediated inhibition is *C. difficile* strain-specific**
A) Colony counts of *Bacteroides* spp. and *C. difficile* 630 from mono- and coculture at 24 h. N = 4, mean ± SD shown. Two-tailed Mann-Whitney or unpaired t-tests were performed to compare each respective monoculture vs coculture counts. **B**) Growth curves of single species as measured by $OD_{600}$ over 12 h. N = 3, mean ± SD is shown. **C**) Growth rates were calculated from growth curve data generated in (**B**) using Growthcurver. Median + interquartile range is shown. A Kruskal-Wallis test was performed, followed by Dunn's correction. ns $P > 0.05$, * $P < 0.05$.

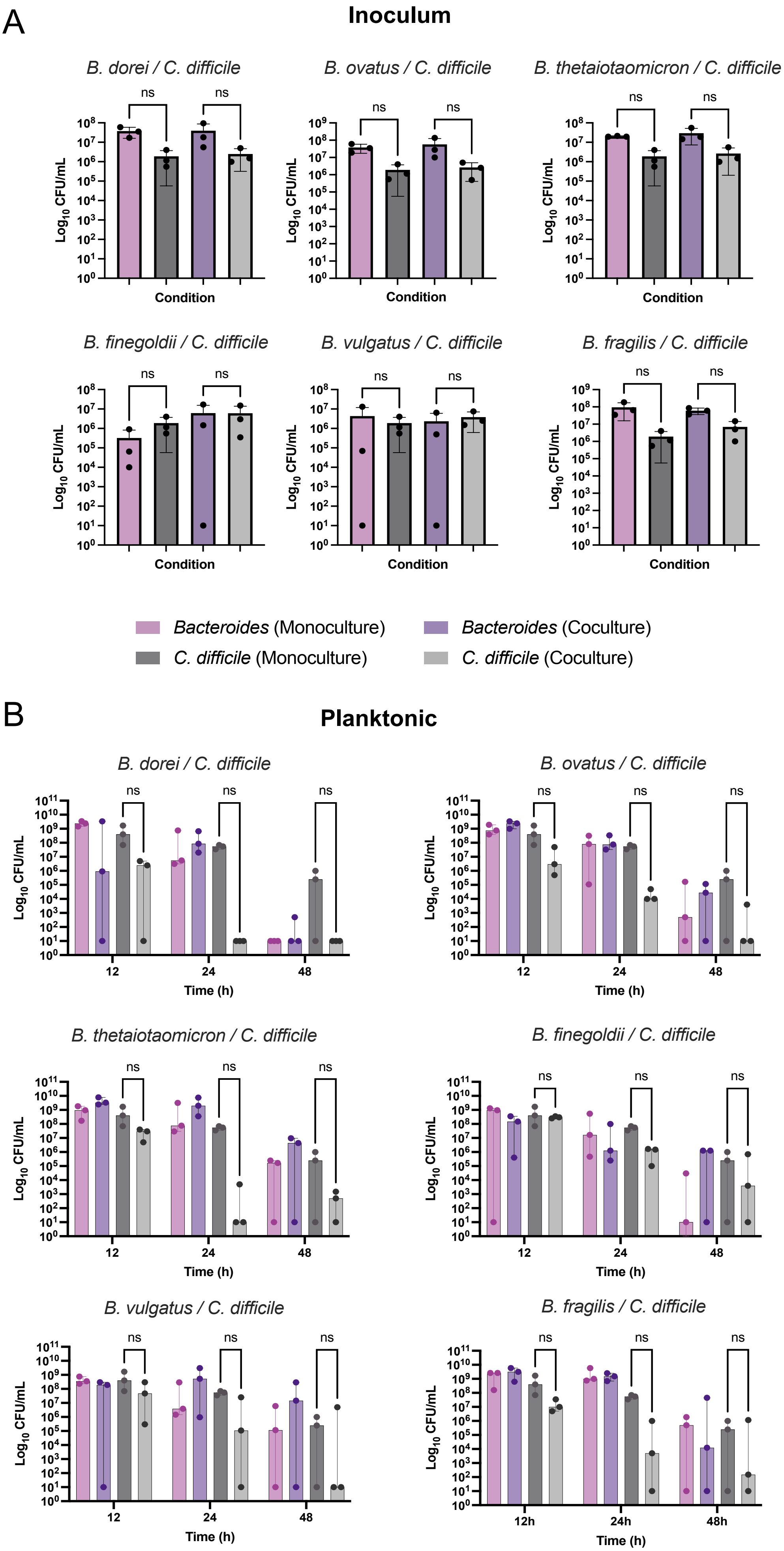
A
Inoculum
B. dorei / C. difficile
B. ovatus / C. difficile
B. thetaiotaomicron / C. difficile
B. finegoldii / C. difficile
B. vulgatus / C. difficile
B. fragilis / C. difficile
ns
Log10 CFU/mL
Condition
Bacteroides (Monoculture)
Bacteroides (Coculture)
C. difficile (Monoculture)
C. difficile (Coculture)
B
Planktonic
B. dorei / C. difficile
B. ovatus / C. difficile
B. thetaiotaomicron / C. difficile
B. finegoldii / C. difficile
B. vulgatus / C. difficile
B. fragilis / C. difficile
ns
Log10 CFU/mL
12
24
48
12h
24h
48h
Time (h)

**Figure S4. *Bacteroides*-mediated inhibition of *C. difficile* is not as potent in planktonic milieu.**
**A**) Bacterial numbers in the inoculum was measured by CFU/mL, from *Bacteroides spp./C. difficile* cocultures and respective single species monocultures. N = 3, mean ± SD is shown. A one-way ANOVA or Kruskal-Wallis test was carried out with Tukey's or Dunn's correction, respectively. **B)** Monoculture and cocultures from A) were grown planktonically over 48 h, and CFU/mL were enumerated. N = 3, median + interquartile range is shown. Where bacterial numbers were below the detection limit, an arbitrary value of 10 was imputed to allow for statistical analysis. Multiple Mann-Whitney tests were performed, comparing each species CFU/mL between coculture and monoculture, at each respective timepoint. ns $P > 0.05$.

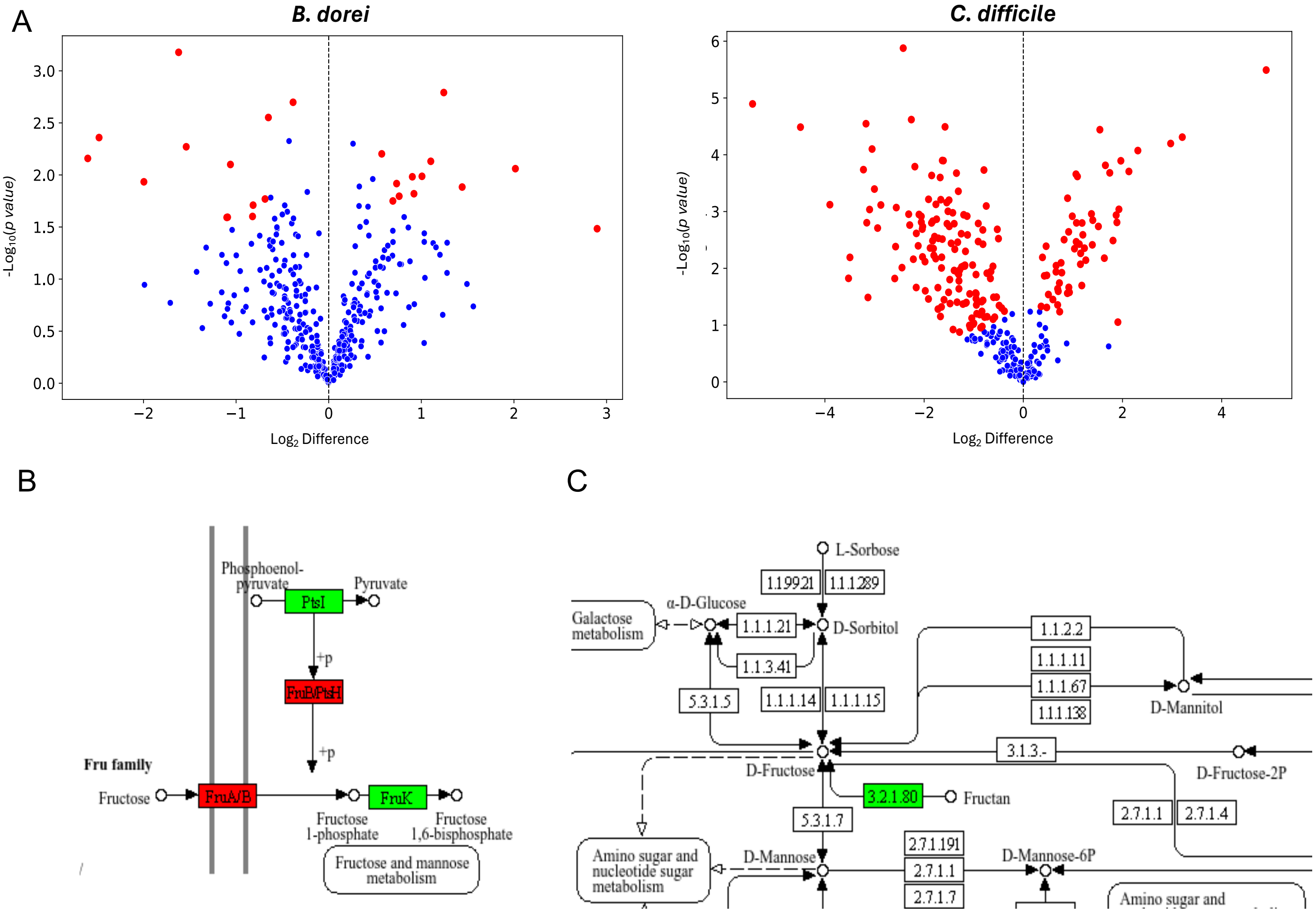


Figure S5. **Mass spectrometry from *B. dorei/C. difficile* biofilm coculture and monoculture biofilm supernatants at 24 h.**
Significantly differentially abundant proteins identified by mass-spectrometry from supernatants of *B. dorei-C. difficile* coculture biofilms after 24 h, relative to monoculture biofilm supernatants. A two-sample t-test was performed. False Discovery Rate (FDR) was set to 0.05 and s0 was set as 0.1 (default for Perseus). Significant genes that passed these cut-offs are coloured in red, with non-significant proteins coloured in blue. **B-C**) KEGG pathways for PTS pathways enriched in *C. difficile* (**B**) and  fructose and mannose metabolism enriched in *B. dorei* in, coculture (**C**). DEPs that were significantly more abundant in the *B. dorei/C. difficile* coculture are coloured in green, those that were significantly more abundant in the monocultures are coloured in red.  N = 3.

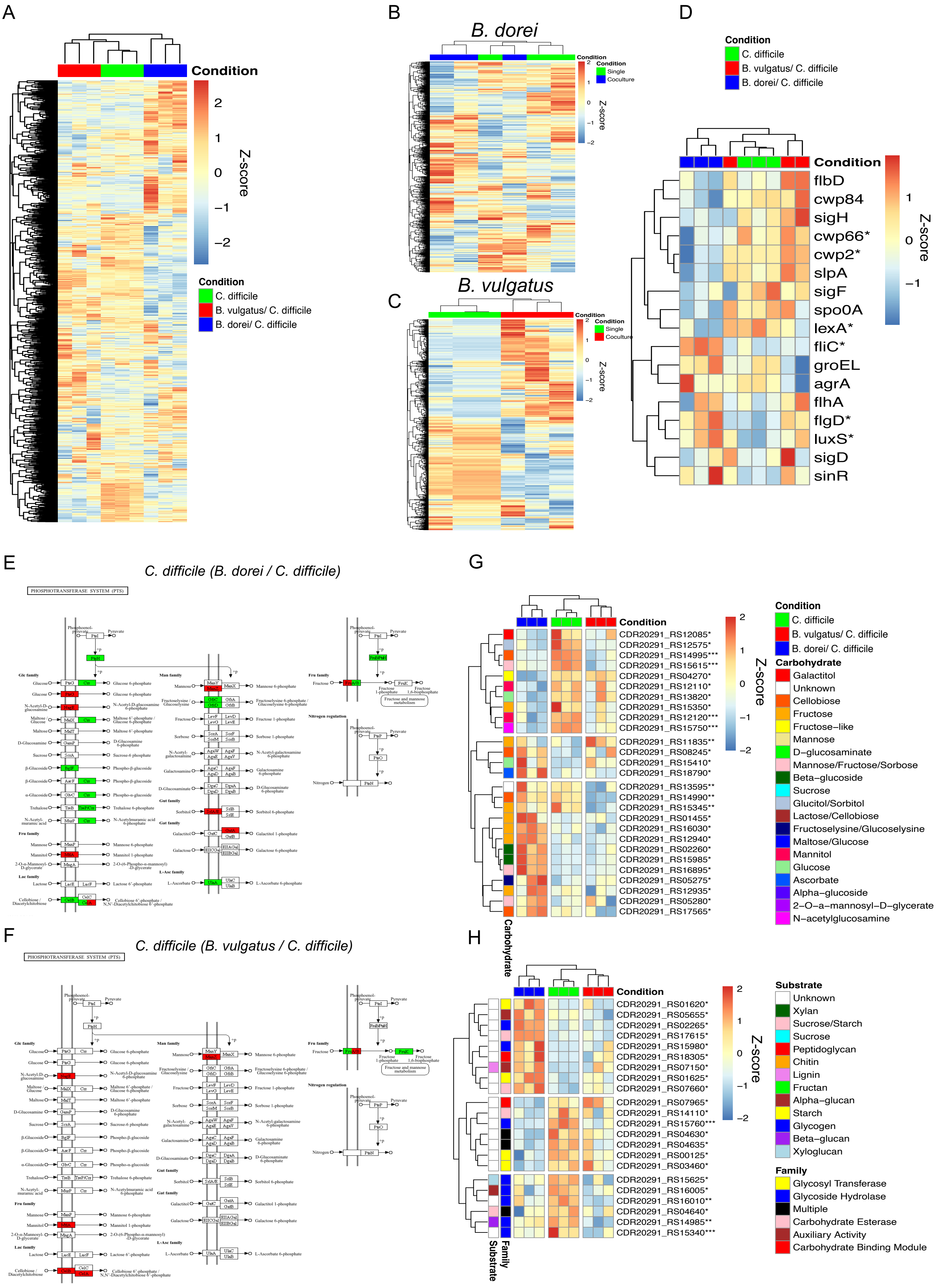

A
Condition
2
1
0
−1
−2
Z-score
Condition
C. difficile
B. vulgatus/ C. difficile
B. dorei/ C. difficile
B
B. dorei
Condition
Condition
Single
Coculture
Z-score
C
B. vulgatus
Condition
Condition
Single
Coculture
Z-score
D
Condition
C. difficile
B. vulgatus/ C. difficile
B. dorei/ C. difficile
Condition
flbD
cwp84
sigH
cwp66*
cwp2*
slpA
sigF
spo0A
lexA*
fliC*
groEL
agrA
flhA
flgD*
luxS*
sigD
sinR
Z-score
E
C. difficile (B. dorei / C. difficile)
PHOSPHOTRANSFERASE SYSTEM (PTS)
F
C. difficile (B. vulgatus / C. difficile)
PHOSPHOTRANSFERASE SYSTEM (PTS)
G
Condition
CDR20291_RS12085*
CDR20291_RS12575*
CDR20291_RS14995***
CDR20291_RS15615***
CDR20291_RS04270*
CDR20291_RS12110*
CDR20291_RS13820*
CDR20291_RS15350*
CDR20291_RS12120***
CDR20291_RS15750***
CDR20291_RS11835**
CDR20291_RS08245*
CDR20291_RS15410*
CDR20291_RS18790*
CDR20291_RS13595**
CDR20291_RS14990**
CDR20291_RS15345**
CDR20291_RS01455*
CDR20291_RS16030*
CDR20291_RS12940*
CDR20291_RS02260*
CDR20291_RS15985*
CDR20291_RS16895*
CDR20291_RS05275*
CDR20291_RS12935*
CDR20291_RS05280*
CDR20291_RS17565*
Carbohydrate
Z-score
Condition
C. difficile
B. vulgatus/ C. difficile
B. dorei/ C. difficile
Carbohydrate
Galactitol
Unknown
Cellobiose
Fructose
Fructose−like
Mannose
D−glucosaminate
Mannose/Fructose/Sorbose
Beta−glucoside
Sucrose
Glucitol/Sorbitol
Lactose/Cellobiose
Fructoselysine/Glucoselysine
Maltose/Glucose
Mannitol
Glucose
Ascorbate
Alpha−glucoside
2−O−a−mannosyl−D−glycerate
N−acetylglucosamine
H
Condition
CDR20291_RS01620*
CDR20291_RS05655*
CDR20291_RS02265*
CDR20291_RS17615*
CDR20291_RS15980*
CDR20291_RS18305*
CDR20291_RS07150*
CDR20291_RS01625*
CDR20291_RS07660*
CDR20291_RS07965*
CDR20291_RS14110*
CDR20291_RS15760***
CDR20291_RS04630*
CDR20291_RS04635*
CDR20291_RS00125*
CDR20291_RS03460*
CDR20291_RS15625*
CDR20291_RS16005*
CDR20291_RS16010**
CDR20291_RS04640*
CDR20291_RS14985**
CDR20291_RS15340***
Family
Substrate
Z-score
Substrate
Unknown
Xylan
Sucrose/Starch
Sucrose
Peptidoglycan
Chitin
Lignin
Fructan
Alpha−glucan
Starch
Glycogen
Beta−glucan
Xyloglucan
Family
Glycosyl Transferase
Glycoside Hydrolase
Multiple
Carbohydrate Esterase
Auxiliary Activity
Carbohydrate Binding Module

I

*B. vulgatus (B. vulgatus / C. difficile)*

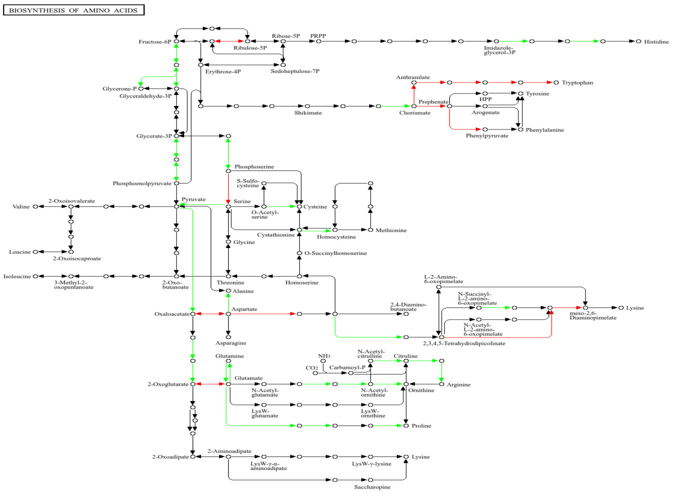

J

*B. dorei (B. dorei / C. difficile)*

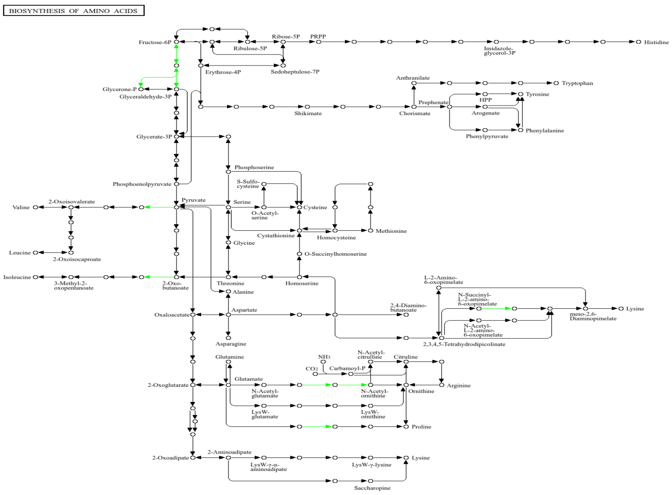

Figure S6. **Heatmaps and pathway maps showing expression profiles from 24 h biofilms**. A-C) Heatmaps showing overall expression profiles from 24 h biofilms from RNAseq data presented as a Z-score for *C. difficile* in different coculture conditions and monoculture control (**A**), and *B. dorei* expression (**B**) or *B. vulgatus* expression (**C**) when cultured on its own or with *C. difficile*. **D**) Expression of *C. difficile* genes associated with biofilm formation within different monoculture and coculture biofilms at 24 h. E-F) KEGG maps of significantly differentially expressed *C. difficile* PTS pathways in *B. dorei* (**E**) and *B. vulgatus* (**F**) coculture biofilms. **G**) Heatmap (presented as a Z-score) showing significant differential expression of *C. difficile* PTS genes in different coculture biofilms, annotated with the respective transported carbohydrate. **H**) Heatmap (presented as a Z-score) of *C. difficile* CAZyme genes differentially expressed in different coculture biofilms, along with their predicted glycan substrates, as predicted by dbCan3**. I-J)** KEGG maps of *B. vulgatus* (**I**) or *B. dorei* (**J**) genes involved in amino acid biosynthetic pathways that are significantly differentially expressed in coculture with *C. difficile* in biofilms. E,F,I,J: significantly upregulated or downregulated genes were coloured in green and red, respectively. For heatmaps A-D,G and H, * denote significant differential expression in *C. difficile* when cocultured with *B. dorei* (*) or *B. vulgatus* (**) relative to the monoculture. *** denotes significant differential expression in both cocultures relative the monoculture condition. $Log_2FC \leq -1$ or $\geq 1$, padj $< 0.05$.

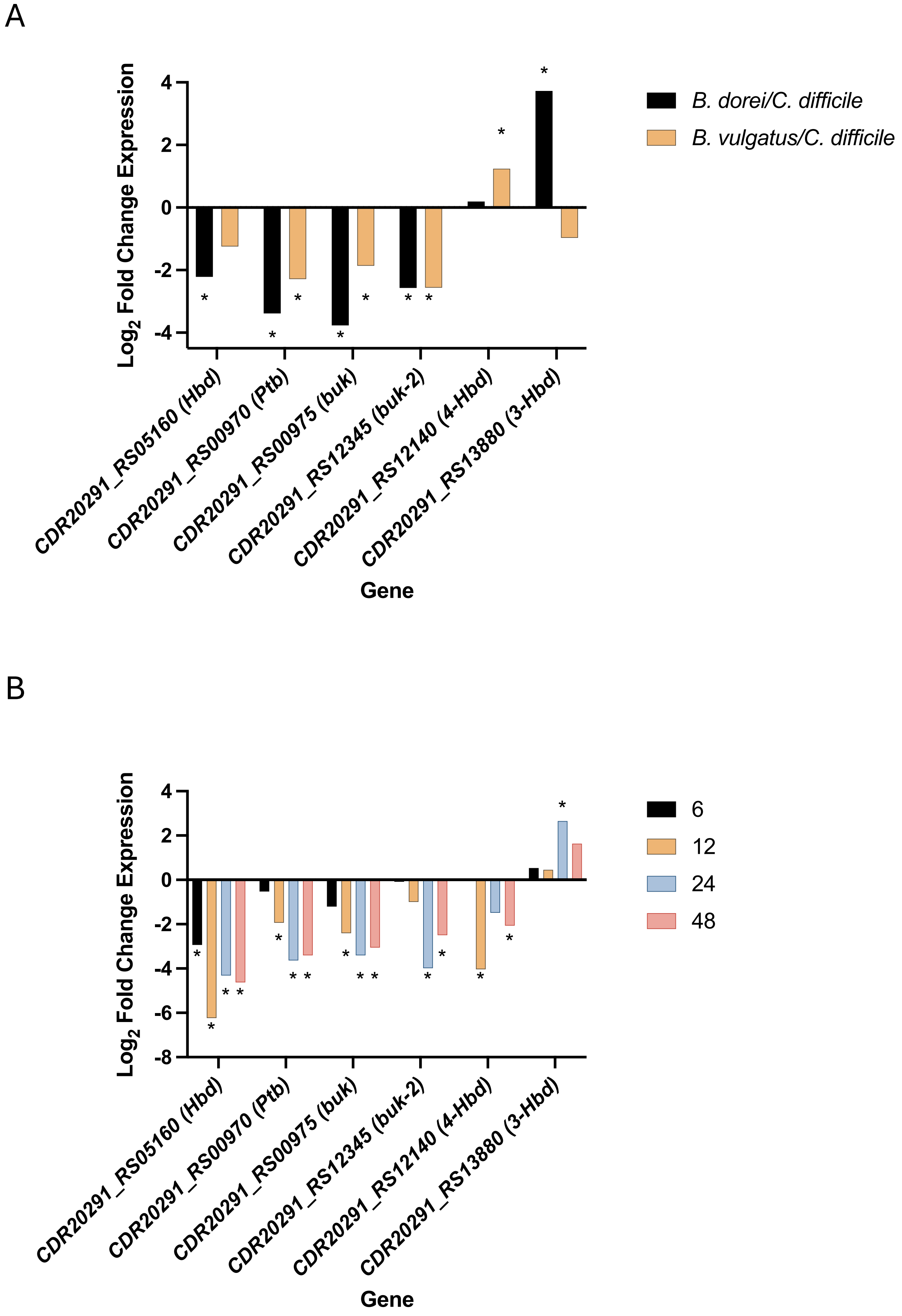


Fig. S7. ***C. difficile* butyrate metabolism is downregulated in a *Bacteroides* and microbiota community cocultures.**

Graphs display the $Log_2FC$ from RNAseq of some *C. difficile* genes involved in butyrate metabolism in either the *Bacteroides* coculture relative to a *C. difficile* monoculture biofilm (**A**), or a commensal community + *C. difficile* biofilm, relative to a control commensal community biofilm at each respective timepoint (**B**). * denotes significant differential expression ($Log_2FC \leq 1$ or $\geq 1$, padj $< 0.05$).

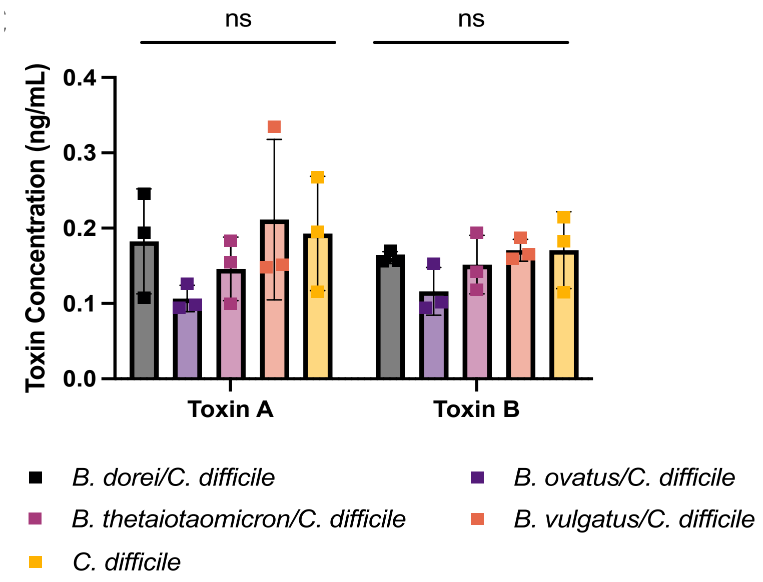


Figure S8. **Toxin levels do not differ in planktonic cocultures.**
Toxin concentrations quantified from planktonic cocultures and *C. difficile* monocultures at 24 h. A Kruskal-Wallis or one-way ANOVA were performed on toxin A ($P = 0.2549$) and toxin B ($P = 1.431$), respectively. N = 3, mean ± SD shown. All samples fell below the detectability cut-off. ns $P > 0.05$.

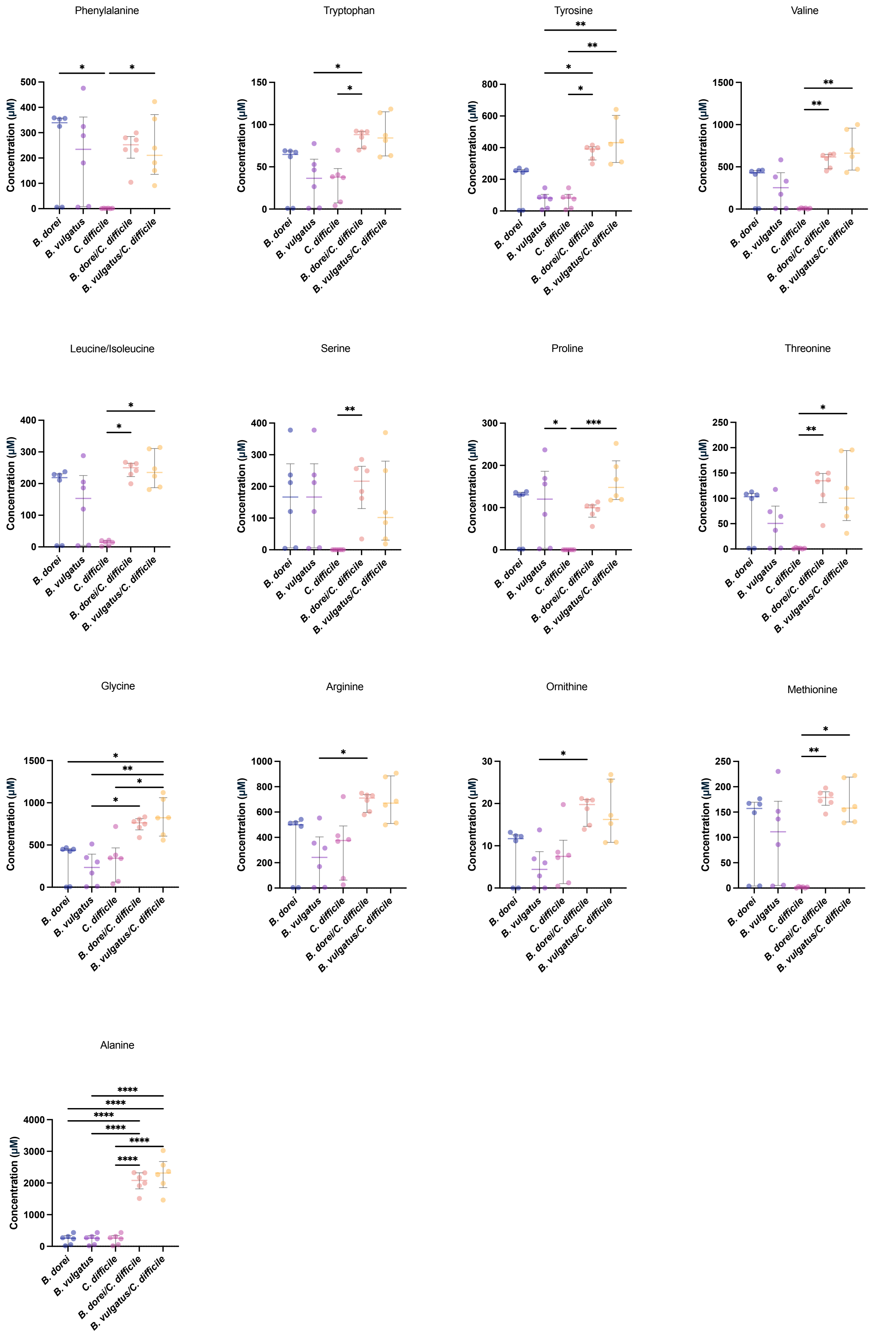
Phenylalanine
Tryptophan
Tyrosine
Valine
Leucine/Isoleucine
Serine
Proline
Threonine
Glycine
Arginine
Ornithine
Methionine
Alanine
Concentration (µM)
B. dorei
B. vulgatus
C. difficile
B. dorei/C. difficile
B. vulgatus/C. difficile
*
**
***
****

Figure S9. **Concentrations of Stickland substrates as quantified by mass spectrometry.** Metabolomic analysis of supernatants from various monoculture and coculture biofilms at 24 h. For concentrations that were NA, an arbitrary value of 0 was assigned to allow for statistical analyses. A Kruskal-Wallis or One-way ANOVA analysis was performed to compare between conditions, followed Dunn's or Tukey's multiple comparison correction, respectively. N = 6. All graphs display the median + interquartile range. * $P < 0.05$, ** $P < 0.01$, *** $P < 0.001$, **** $P < 0.0001$.

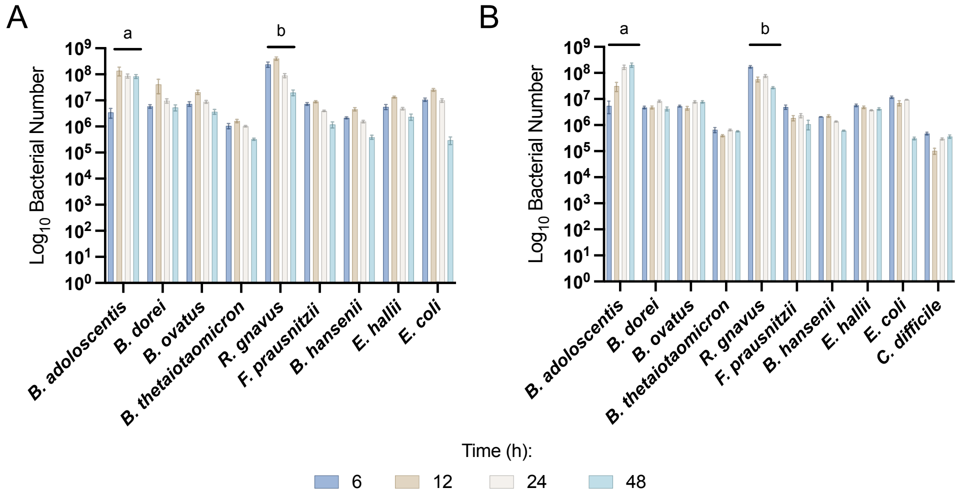


C

| | | Microbiota | | Microbiota + *C. difficile* | |
|---|---|---|---|---|---|
| | | Summary | Adjusted p value | Summary | Adjusted *p* value |
| *B. adolescentis* | ***a*** | | | | |
| | *6 vs. 12* | **** | <0.0001 | ** | 0.0038 |
| | *6 vs. 24* | **** | <0.0001 | **** | <0.0001 |
| | *6 vs. 48* | **** | <0.0001 | **** | <0.0001 |
| | *12 vs. 24* | ** | 0.0025 | **** | <0.0001 |
| | *12 vs. 48* | ** | 0.0012 | **** | <0.0001 |
| | *24 vs. 48* | ns | 0.9965 | *** | 0.0001 |
| *R. gnavus* | ***b*** | | | | |
| | *6 vs. 12* | **** | <0.0001 | **** | <0.0001 |
| | *6 vs. 24* | **** | <0.0001 | **** | <0.0001 |
| | *6 vs. 48* | **** | <0.0001 | **** | <0.0001 |
| | *12 vs. 24* | **** | <0.0001 | * | 0.0466 |
| | *12 vs. 48* | **** | <0.0001 | *** | 0.0007 |
| | *24 vs. 48* | **** | <0.0001 | **** | <0.0001 |

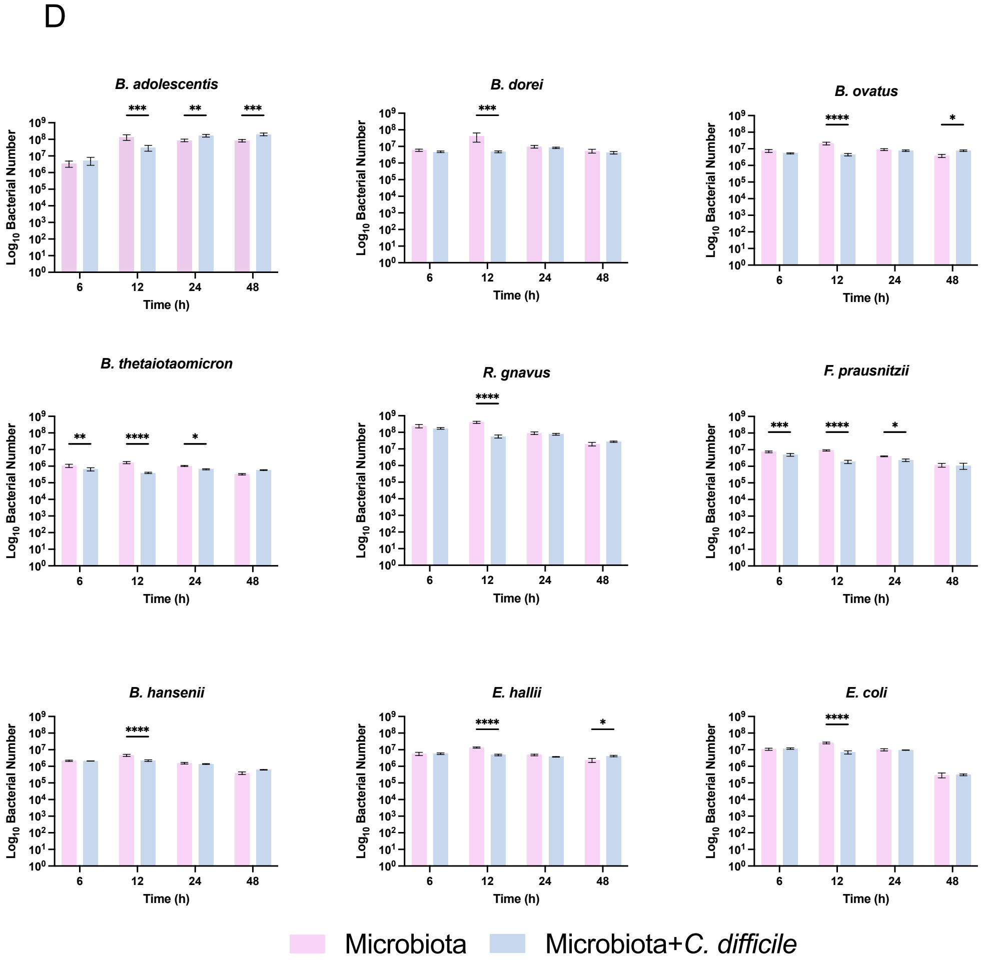

Figure S10. **Bacterial numbers of individual species within the commensal community ± *C. difficile***

Bacterial numbers within multispecies biofilms cultured in presence or absence of *C. difficile* were calculated using PMA-qPCR at 6-48 h. A-B represent the calculated number of bacteria per species over time in the microbiota (**A**) and microbiota + *C. difficile* community (**B**) as deduced by PMA-qPCR. A two-way ANOVA was performed on each, followed by a Tukey's multiple comparison correction. The a and b denotations represent significance which can be found in panel C. **D**) Bacterial numbers of individual species within the commensal community ± *C. difficile* over time were compared as deduced by PMA-qPCR. Two-way ANOVAs were performed. Šídák correction was performed for comparisons between condition at each respective timepoint. All data presented is of three biological replicates, each with three technical replicates, mean ± SD is shown. ns $P > 0.05$, * $P < 0.05$, ** $P < 0.01$, *** $P < 0.001$, **** $P < 0.0001$.

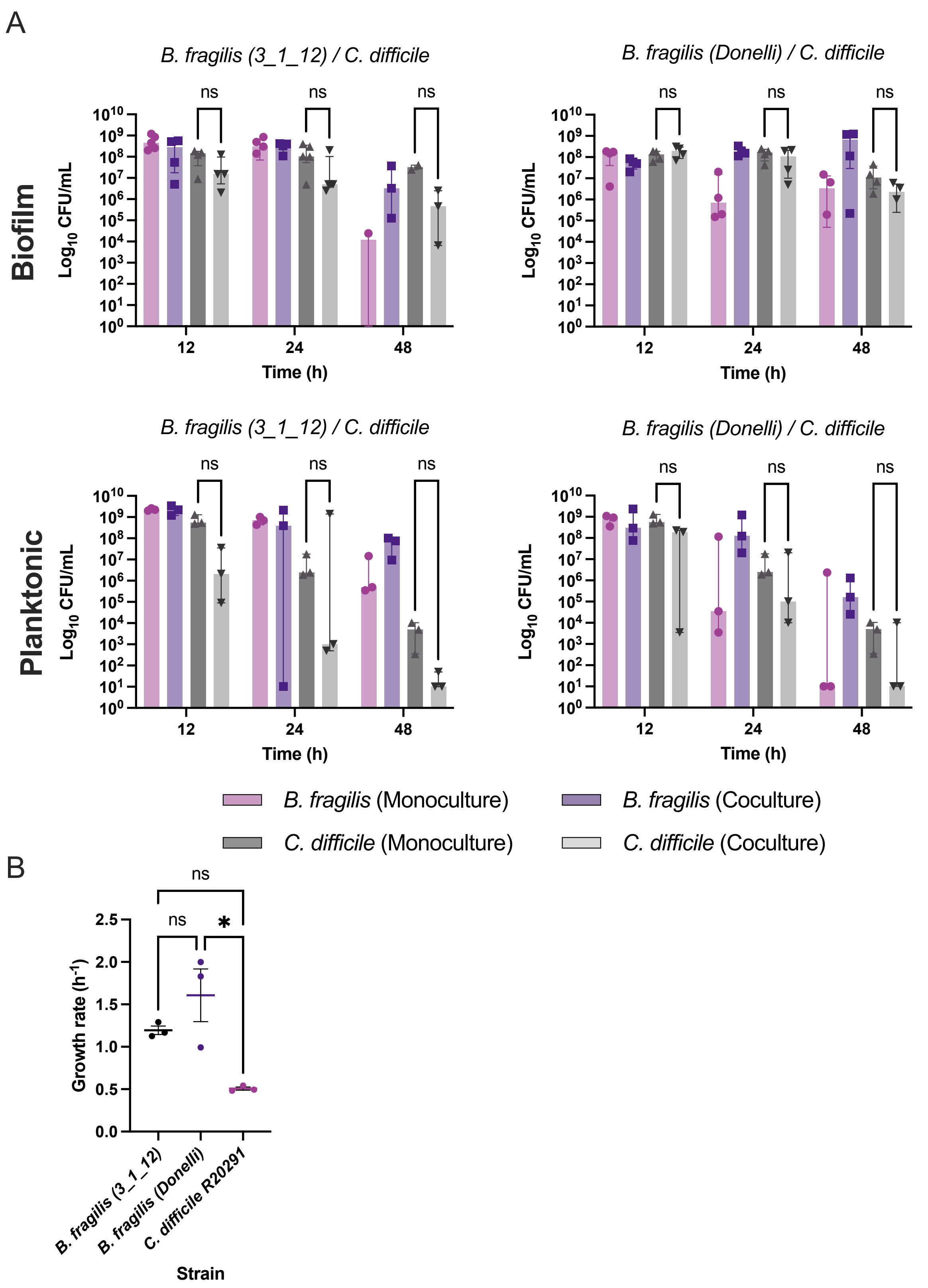


Figure S11. **Comparison of two strains of *B. fragilis* with *C. difficile* in biofilm and planktonic coculture.**

**A)** *B. fragilis* strain 3_1_12 or a clinical isolate[39] were cultured with *C. difficile* in biofilms or planktonic culture over 48 h, as measured by CFU/ml. Data represents a minimum of three biological repeats. Median plus interquartile range is shown. Multiple unpaired t-tests showed no significant differences. **B)** Planktonic growth curves over 12 h were performed and growth rates were calculated using Growthcurver. N = 3, mean ± SD is shown. A one-way ANOVA followed by Dunnett's multiple comparison test was performed. ns $P < 0.05$, * $P < 0.05$.
