## Supplemental Methods for "*Bacteroides*-driven metabolic remodelling suppresses *Clostridioides difficile* toxin expression in mixed biofilm communities"

### Proteomics

Supernatants from biofilms were retrieved at the desired timepoint, centrifuged 1500 x *g*, 5 min, filter sterilised through a 0.2 µm filter, and treated with 1X EDTA-free protease inhibitor (Roche, Sigma-Aldrich, USA). Supernatants were prepped for mass spectrometry using the Filter Aided Sample Preparation (FASP) method. Unless stated otherwise, centrifuge steps were performed at RT for 20 min at 8000 x *g.* 200 µg of sample was applied to a spin column, and centrifuged. A 400 µL aliquot of 50 mM ammonium bicarbonate was added, and each tube was spun. This was repeated an additional 4 times. After the final ammonium bicarbonate wash, the supernatant was discarded, and the samples were reduced and alkylated in 400 µL ammonium bicarbonate + 10 mM Tris(2-carboxyethyl)-phosphine (TCEP) (Sigma-Aldrich, USA) and 40 mM 2-chloroacetamide (CAA) (Sigma-Aldrich, USA) at RT for 30 min. Columns were centrifuged and subsequently washed 5 times with 400 µL 50 mM ammonium bicarbonate. After discarding the final wash supernatant, 4 µg trypsin (Promega) was added and made up to 400 µL with ammonium bicarbonate and the sample was incubated overnight at 37 °C. Following overnight incubation, the columns were moved to new collection tubes and the samples were centrifuged, followed by the addition of 400 µL MS-water and an additional spin. Samples were then centrifuged in a SpeedVac (Eppendorf, Germany ) for approximately 2.5 hr at 50 °C, and the samples were subsequently resuspended in 50 µL 2% acetonitrile (CAN) 0.1% trifluoroacetic acid (TFA).

The resuspended samples were moved to a HPLC tube and mass spectrometry, and subsequent preliminary data filtering and analysis was carried out by Dr. Cleidiane Zampronio (WPH Proteomics Facility RTP). Briefly, this involved injecting 1 µL of each sample into a Q-Exactive LC-MS Orbitrap Fusion (Thermo Fisher Scientific, USA) for proteomic analysis. Here, reversed phase chromatography was used to separate tryptic peptides before mass spectrometry analysis. The Ultimate 3000 RSLCnano system (Thermo Fisher Scientific, USA) was used with mobile phase buffer A (0.1 % formic acid in water), and mobile buffer B (0.1 % formic acid in CAN). 1 µL of each sample was loaded onto a µ-precolumn cartridge (Thermo Fisher Scientific, USA) equilibrated in 2 % aqueous CAN containing 0.1 % TFA and peptides were subsequently eluted onto an analytical column (Bruker nanoElute Forty Analytical column) at 350 nL $min^{-1}$. This was done by incrementally increasing the mobile B concentration from 4 % to 25 % over 36 min, increasing it to 35 % over 10 min, and finally to 90 % over 3 min, followed by a 10 min re-equilibration at 4 % mobile B concentration. The Ultimate 3000 RSLCnano system was coupled online to a hybrid timsTOF pro (Bruker Daltonics, USA) using a CaptiveSpray nano-electrospray ion source operated in Data-Dependent Parallel Accumulation-Serial Fragmentation (PASEF) mode. Peptide separation was done based on ion mobility depending on their collisional cross sections and charge states. Method settings used were set at: mass range 100-1700 m/z, ion mobility range 1/K0 Start 0.6 $Vs/cm^2$ End 1.6 Vs/cm2, Ramp rate 9.42 Hz and Duty cycle 100 %. Raw mass spectrometry data was searched against the Uniprot proteome databases UP000002070 for *C. difficile* and UP000293499 for *B. dorei*, using FragPipe (Version 18.0) [90], and a common contaminant database. For this database search, peptides were generated from a tryptic digestion allowing for up to two missed cleavages, with carbamidomethylation of cysteine residues as fixed modifications. Oxidation of methionine and acetylation of the protein N-terminus were added as variable modifications. Mass spectrometry data was mapped to respective species proteomes and was analysed using Perseus (Version 2.0.5.0) and preliminarily viewed on Scaffold (Version 5.3.3). From the LFQ intensity output, the data was filtered, removing low probability (>0.1) entries. Data was subsequently transformed to take the $Log_2$ value, grouped by conditions, and then high stringent filtering was performed, removing proteins that did not appear in all three replicates in at least one condition. Missing values were then imputed to allow for normal distribution. Downstream analysis was carried out using Perseus' in-built statistical tests and plotting functions.
