## Supplemental Tables S1-S3 for "*Bacteroides*-driven metabolic remodelling suppresses *Clostridioides difficile* toxin expression in mixed biofilm communities"

**Table S1 List of bacterial species and strains**

| Species | Phylum | Source |
|---|---|---|
| *Clostridioides difficile (R20291)* | Firmicutes | Stoke Mandeville; Trevor Lawley |
| *Clostridioides difficile (630)* | Firmicutes | Trevor Lawley |
| *Bifidobacterium adolescentis (L2-32)* | Actinobacteria | BEI Resources |
| *Bacteroides dorei (CL02T00C15)* | Bacteroidetes | BEI Resources |
| *Bacteroides thetaiotaomicron (3_8_47FAA)* | Bacteroidetes | Anne Marie Krachler |
| *Ruminococcus gnavus (VPI-5482)* | Firmicutes | BEI Resources |
| *Faecalibacterium prausnitzii (CC55_001C)* | Firmicutes | DSMZ |
| *Blautia hansenii (17677 A2-165)* | Firmicutes | DSMZ |
| *Eubacterium hallii (3353)* | Firmicutes | DSMZ |
| *Escherichia coli (83972)* | Proteobacteria | BEI Resources |
| *Bacteroides vulgatus (CL09T03C04)* | Bacteroidetes | BEI Resources |
| *Bacteroides finegoldii* (CL09T03C10) | Bacteroidetes | BEI Resources |
| *Bacteroides fragilis (3_1_12)* | Bacteroidetes | BEI Resources |
| *Bacteroides fragilis* | Bacteroidetes | Gianfranco Donelli |

**Table S2 List of primers used for RT-qPCR and PMA-qPCR**

| Species | Forward primer (5' to 3') | Reverse primer (5' to 3') | Gene target |
|---|---|---|---|
| **RT-qPCR** | | | |
| *C. difficile* | TCGTGGGAATTTAGCTGCAG | TTCCCAACGGTCTAGTCCAA | *tcdA* |
| *C. difficile* | AGGAGGCGTTATGAATATGACA | TGCTACTTTTCTGATTCCTCCA | *tcdE* |
| *C. difficile* | AACTCAGTAGATGATTTGCAAGAA | TCTCCCTCTTCATAATGTAAAACTC | *tcdR* |
| **PMA-qPCR** | | | |
| *C. difficile* | GGTTGAAAGAATAGCAGAGTTAGTT | GCATTAGCATCCCTCTTTAATTCTA | *gyrA* |
| *B. adolescentis* | CTCCGGATACACGGTCATGG | GTCTTCGATATCCACGCCGA | *topI* |
| *B. dorei* | AAGCGGCTTCAAGAAACAGG | GTGCCCTTTACCTTGGGAAC | *topI* |
| *B. ovatus* | GGGCCTATTATCGCAACCGA | AGGTGCATACGTAGACGGAC | *topI* |
| *B. thetaiotaomicron* | GTCTGTAATCAAGTCCGCCG | AATGCCGGAAAGCGGTAAAC | *topI* |
| *R. gnavus* | GCTGAACAGAGCAGAAGAGC | TCCTTCGCAGTCTGAACATTCT | *gyrA* |
| *F. prausnitzii* | CCGGTGTCCGTGTCATGC | CTCAGCCTCTACTGTCTCGG | *gyrA* |
| *B. hansenii* | GACGTAAGAAGCACCGGTAGA | ATAATCGCCCTGACAGGTAAGC | *gyrA* |

| | | | |
|---|---|---|---|
| *E. hallii* | TACCGCCTCATCGGACTTGA | TCATGGAGGCTGGATGCTCT | *gyrA* |
| *E. coli* | GAACTCGGTGAGGACGGTTT | GCTGGAACAGGACGAACGTA | *gyrA* |

**Table S3. Accession numbers used in this study.**

| **RNA sequencing analysis** | |
|---|---|
| **Species** | **NCBI Accession Number** |
| *B. adolescentis* | GCF_000154085.1 |
| *B. dorei* | GCF_000273035.1 |
| *B. ovatus* | GCF_000218325.1 |
| *B. thetaiotaomicron* | GCF_000011065.1 |
| *R. gnavus* | GCF_000507805.1 |
| *F. prausnitzii* | GCF_010509575.1 |
| *E. Halllii* | GCF_000173975.1 |
| *E. coli* | GCF_000148365.1 |
| *C. difficile R20291* | GCF_000027105.1 |
| *B. vulgatus* | GCF_000273295.1 |
| **Proteomics analysis** | |
| ***Species*** | **Uniprot Accession Number** |
| *B. dorei* | UP000293499 |
| *C. difficile* | UP000002070 |
